# Structural insights into the allosteric inhibition of P2X4 receptors

**DOI:** 10.1101/2023.03.07.531612

**Authors:** Cheng Shen, WenWen Cui, Yuqing Zhang, Yimeng Zhao, Danqi Sheng, Xinyu Teng, Jin Wang, Motoyuki Hattori

## Abstract

P2X receptors are ATP-activated cation channels involved in a variety of physiological functions. Among the seven subtypes of P2X receptors, the P2X4 subtype plays important roles in both the immune system and the central nervous system, particularly in neuropathic pain. Therefore, P2X4 receptors are of increasing interest as potential drug targets, and several P2X4 subtype-specific inhibitors have been developed. However, the mechanism of allosteric inhibition of P2X4 receptors remains largely unclear due to the lack of structural information.

Here, we report the cryo-EM structures of the zebrafish P2X4 receptor in complex with two P2X4 subtype-specific antagonists, BX430 and BAY-1797. Both antagonists bind to the same allosteric site located at the subunit interface at the top of the extracellular domain. Structure-based mutational analysis by electrophysiology identified the important residues for the allosteric inhibition of both zebrafish and human P2X4 receptors.

Interestingly, in the previously reported apo structure, the binding pocket is closed and too narrow to accommodate allosteric modulators. Structural comparison revealed the ligand-dependent structural rearrangement of the binding pocket to stabilize the binding of allosteric modulators, which in turn would prevent the structural changes of the extracellular domain associated with channel activation. Furthermore, comparison with the previously reported P2X structures of other subtypes provided mechanistic insights into subtype-specific allosteric inhibition.

Overall, the present work provides structural insights into the allosteric inhibition mechanism of P2X4 receptors, facilitating the design and optimization of specific modulators for P2X4 receptors.

## Introduction

ATP not only is the primary source of energy in cells but also acts as an extracellular signaling molecule^1^. P2X receptors are ATP-activated, nonselective cation channels with diverse biological functions^2–4^. The P2X receptor family consists of seven subtypes, designated P2X1-P2X7, that form homo- or heterotrimers^5, 6^. While the transmembrane (TM) domain of P2X receptors is responsible for ion permeation, the extracellular domain contains an agonist binding site as well as binding sites for regulatory factors, including divalent cations and chemical compounds^7^. With ubiquitous expression patterns, P2X receptors regulate many essential cellular functions, including synaptic transmission, pain perception, inflammatory response, and smooth muscle contraction^8,9,10^.

Among the seven P2X subtypes, the P2X4 receptor subtype is highly Ca^2+^ permeable and is expressed on a variety of cells, including central neurons and immune cells^11, 12^. In particular, P2X4 receptors are expressed in macrophages, microglia and monocytes, where they play a role in the release of proinflammatory cytokines; thus, P2X4 receptors are a promising target for the treatment of neuropathic pain^13–15^.

As increasing evidence highlights the role of P2X4 receptors in physiological and pathological processes, several subtype-specific antagonists have been identified^16^, including BX430^17^ and BAY-1797^18^, and some have entered clinical trials for the treatment of neuropathic pain^8^.

BX430, or 1-(2,6-dibromo-4-isopropylphenyl)-3-(3-pyridyl)urea, is a phenylurea antagonist. BX430 inhibits human P2X4 currents with an IC_50_ value of 0.54 μM and has no significant effect on other subtypes^17^. In addition, BX430 inhibits both human and zebrafish P2X4 but not rat and mouse P2X4^19^. BAY-1797, or N-[4-(3-chlorophenoxy)-3-sulfamoylphenyl]-2-phenylacetamide, is a sulfonamide antagonist of P2X4 with an IC_50_ of 211 nM and high subtype specificity^18^. It shows anti-inflammatory and antinociceptive effects in a mouse model^18^.

In efforts to understand the molecular mechanism of P2X receptors, various structures of P2X receptors of different subtypes (P2X3, P2X4 and P2X7) in different functional states (apo-, agonist- and antagonist-bound states) have been reported^20–29^. For the P2X4 receptor, both apo- and ATP-bound structures of the zebrafish P2X4 receptor have been reported^20, 21^, revealing the channel activation mechanism. In addition, structure determination of P2X3 and P2X7 receptors in complex with antagonists, including negative allosteric modulators, provided the structural basis for the inhibition mechanism of these receptors^23, 24, 26, 29^. However, despite the physiological and pharmacological importance of P2X4 receptors, the detailed mechanism of P2X4 inhibition by antagonists, such as the mechanism of P2X4 subtype-specific allosteric inhibition, remains unclear due to the lack of structures of the P2X4 receptor in complex with its selective antagonists.

Here, we report the cryo-EM structures of the zebrafish P2X4 receptor in complex with two different P2X4-selective antagonists, BX430 and BAY-1797. By combining these structures with structure-based patch-clamp analysis, we characterized an allosteric site at the subunit interface in the extracellular domain of the P2X4 receptor. Structural comparison with the previously determined apo structure and MD simulation revealed the structural rearrangement at the binding pocket of the allosteric site to stabilize allosteric modulator binding, which is associated with channel inactivation. Further comparison with previously determined P2X structures of other subtypes provided mechanistic insights into subtype-specific inhibition. These results will facilitate the structure-guided design and optimization of allosteric modulators for the P2X4 receptor.

## Results

### Functional characterization and structure determination

We first evaluated the effects of BX430 and BAY-1797 by whole-cell patch-clamp recording of the zebrafish P2X4 (zfP2X4) receptor expressed in HEK293 cells (**Supplementary Fig. 1)**. BX430 inhibited ATP-dependent currents of both zebrafish P2X4 and human P2X4 (hP2X4) with comparable IC_50_ values at sub-micromolar levels (**Supplementary Fig. 1A, 1B)**, and the results were consistent with the previously reported IC_50_ value of 0.54 µM for hP2X4^17^. While BAY-1797 has been shown to inhibit the hP2X4 receptor^18^, it was not clear whether BAY-1797 could act at the zfP2X4 receptor. In our experiments, the application of 5 µM BAY-1797 strongly suppressed the ATP-evoked currents from zfP2X4 expressed in HEK293 cells (**Supplementary Fig. 1C, 1D)**.

Next, we used single-particle cryo-EM to determine the structures of the zebrafish P2X4 receptor in complex with BX430 and BAY-1797. The purified zfP2X4 protein was reconstituted into PMAL-C8 amphipol and incubated with 50 μM BX430 or 125 μM BAY-1797 for single-particle cryo-EM analysis. After cryo-EM data acquisition and image processing, the final EM map resolutions were 3.4 Å resolution for the BX430-bound structure and 3.8 Å resolution for the BAY-1797-bound structure (**Supplementary Figs. 2, 3, 4, 5)**.

### Allosteric binding site

The overall structures of the zebrafish P2X4 receptor in complex with BX430 and BAY-1797 are very similar (**Supplementary Fig. 6A)** and show a chalice-like trimeric architecture with each subunit containing a relatively large extracellular domain and two TM helices (**Fig. 1**), resembling the shape of a dolphin (**Supplementary Fig. 7A)**, as in the case of previously reported P2X receptor structures^30^. Consistent with the function of BX430 and BAY-1797 as antagonists, the TM helices of both structures adopt a closed conformation of the channel (**Supplementary Fig. 6B, 6C).**

**Figure 1.**
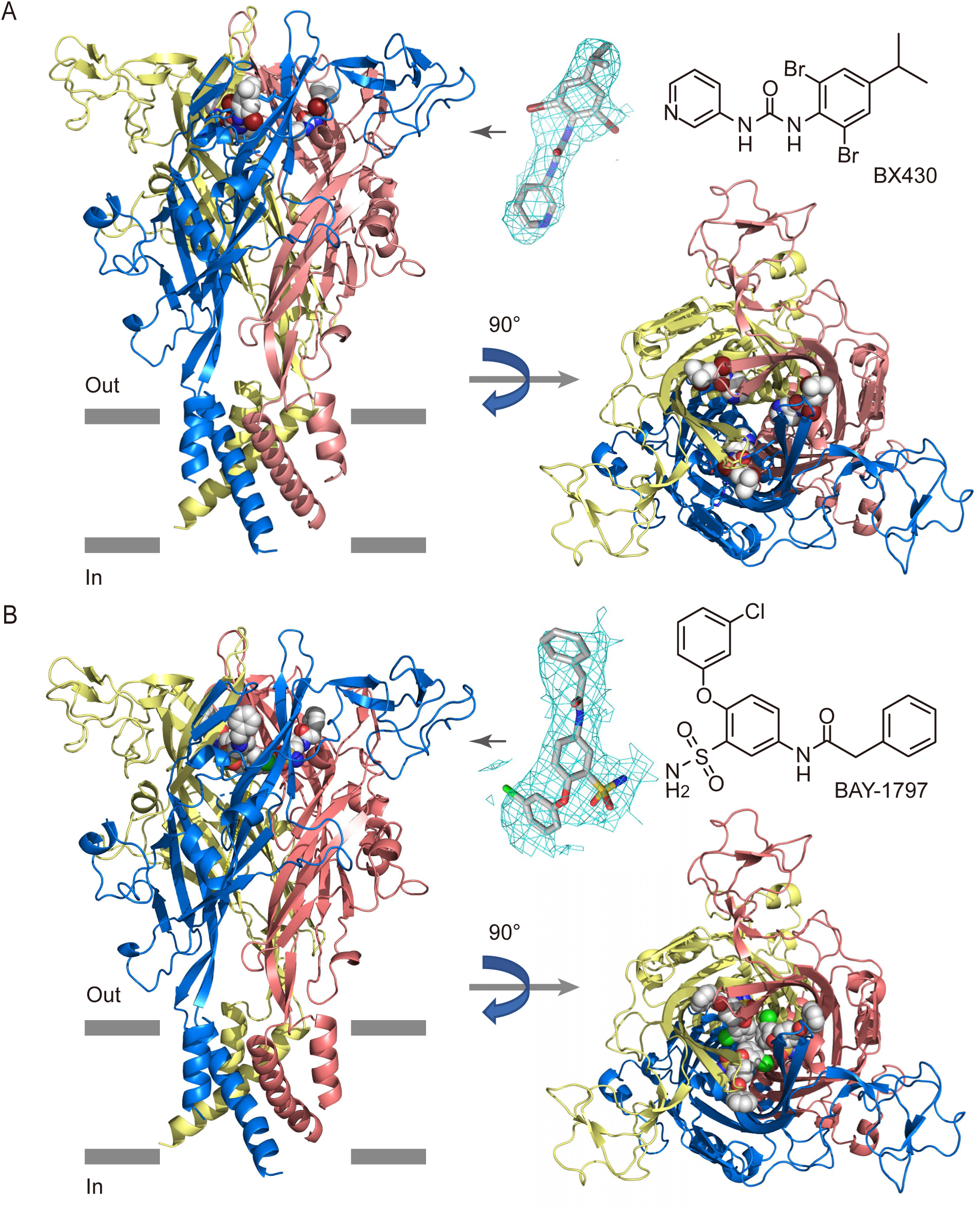
Overall structures. The cryo-EM structures of the zebrafish P2X4 receptor in complex with BX430 (**A**) and BAY-1797 (**B**), viewed parallel to the membrane (left panel) and perpendicular to the membrane from the extracellular side (right panel). BX430 and BAY-1797 molecules bound to the P2X4 receptor are shown in sphere representations. 2D structures of BX430 and BAY-1797 are also shown. The protomers of the trimer are colored blue, yellow, and red. The EM density maps for BX430 and BAY-1797 are also shown and contoured at 4.0 σ and 3.0 σ, respectively.

In the extracellular domain of the two structures, we observed residual EM densities that matched the shapes of BX430 and BAY-1797 at the same location of the intersubunit interface of the trimer (**Fig. 1**). In the dolphin model, this binding site is located in the upper body domain and is distant from the ATP binding site, defining the binding site as an allosteric site (**Supplementary Fig. 7A, 7B)**. To verify the P2X4 structures in complex with BX430 and BAY-1797, we performed MD simulations. During the MD simulations, both overall structures were mostly stable (**Supplementary Fig. 8A, 8C**), and both BX430 and BAY-1797 were stably bound to the allosteric site (**Supplementary Fig. 9**).

In the BX430-bound structure, the dibromo-isopropylphenyl group of BX430 faces the outer side of the receptor, while the pyridine group faces the inner side (**Fig. 2**). The binding site consists mainly of hydrophobic residues, including Trp87, Ile94, Met108, Ile110, Phe299 and Ile315 from one subunit and Tyr302 from the neighboring subunit (**Fig. 2**). In addition, the main chain carbonyl group of Asp91 forms hydrogen bonds with the amines in the urea group of BX430 (**Fig. 2B**). Interestingly, the side chain of Lys301 forms a cation-pi interaction with the aromatic ring of Tyr302 from the neighboring subunit, allowing hydrophobic interactions between the side chain of Lys301 and the 4-isopropylphenyl ring of BX430 (**Fig. 2B**).

**Figure 2.**
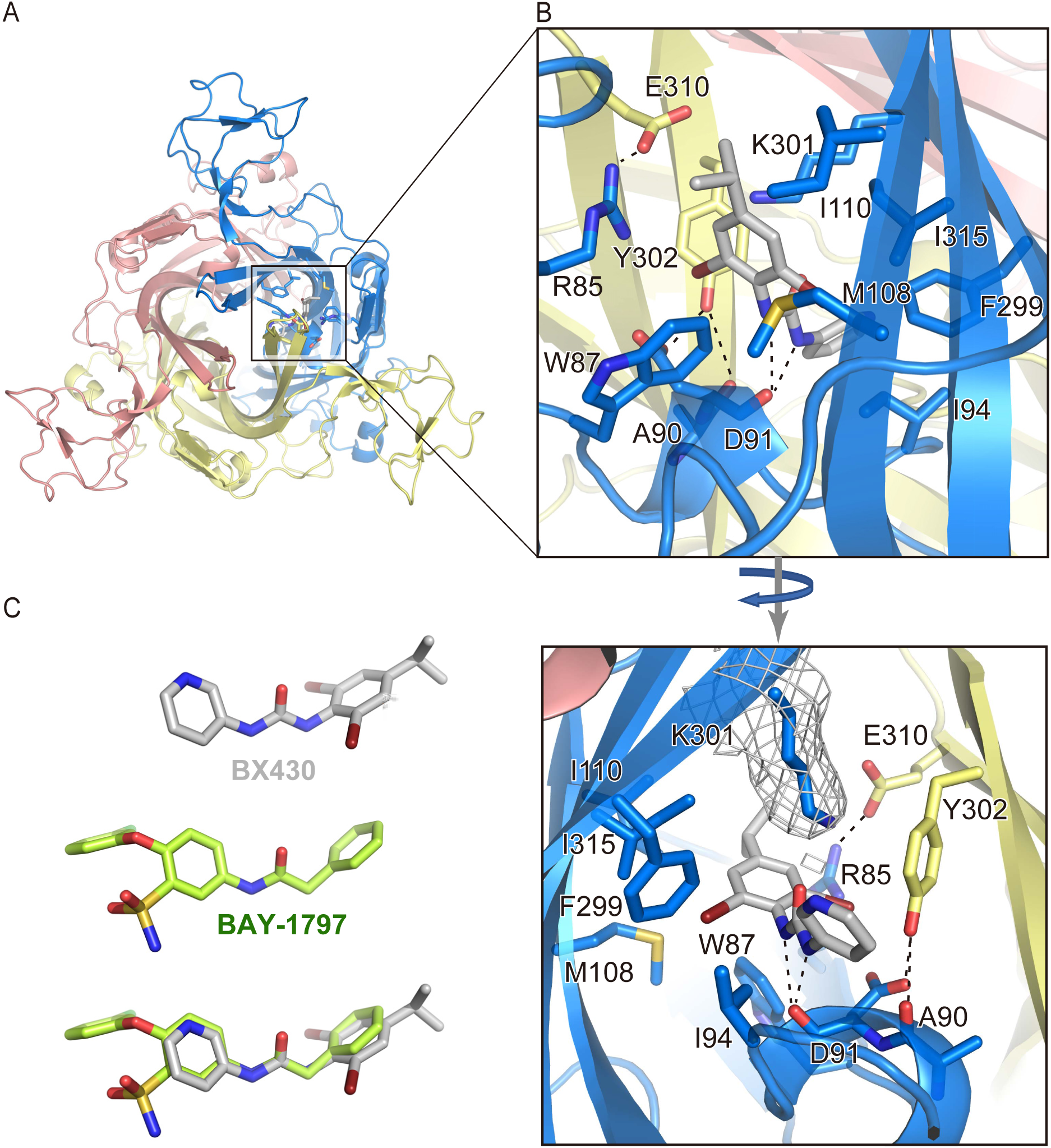
BX430 binding pocket. (**A**) Overall structure of the BX430-bound P2X4 receptor viewed from the extracellular side. The BX430 molecule and amino acid residues involved in its binding from one of three equivalent binding pockets are shown in stick representations. (**B**) Close-up views of the BX430 binding pocket. Dotted lines represent hydrogen bonds. The EM density map for the Lys301 residue is shown and contoured at 3.0 σ. (**C**) Close-up of views of BX430 and BAY-1797 molecules in the cryo-EM structure and the superposition of BAY-1797 onto BX430.

Furthermore, there are intersubunit salt bridges between the side chains of Arg85 of one subunit and Glu310 of the neighboring subunit, and hydrogen bonds between the side chain of Tyr302 of one subunit and the main-chain carbonyl group of Ala90 and the side chain of Asp91 of the neighboring subunit (**Fig. 2B**). These interactions also seemingly stabilize the formation of the allosteric binding pocket.

In the BAY-1797-bound structure, the phenylacetamide moiety faces the outer side of the receptor, while the chlorophenoxy moiety faces the inner side (**Fig. 3**). The BAY-1797 binding site also consists primarily of hydrophobic residues (**Fig. 3B**), which are the same as those of the BX430 binding site (**Fig. 2B**). Furthermore, both Asp91 and Lys301 are similarly involved in BAY-1797 binding (**Fig. 3B**), despite the structural differences between BAY-1797 and BX430. The main chain carbonyl group of Asp91 forms a hydrogen bond with the amine in the phenylacetamide moiety of BAY-1797, and the hydrophobic part of the Lys301 side chain interacts with the phenyl group of the phenylacetamide moiety by forming a cation-pi interaction between the side chain of Lys301 and the aromatic ring of Tyr302 (**Fig. 3B**).

**Figure 3.**
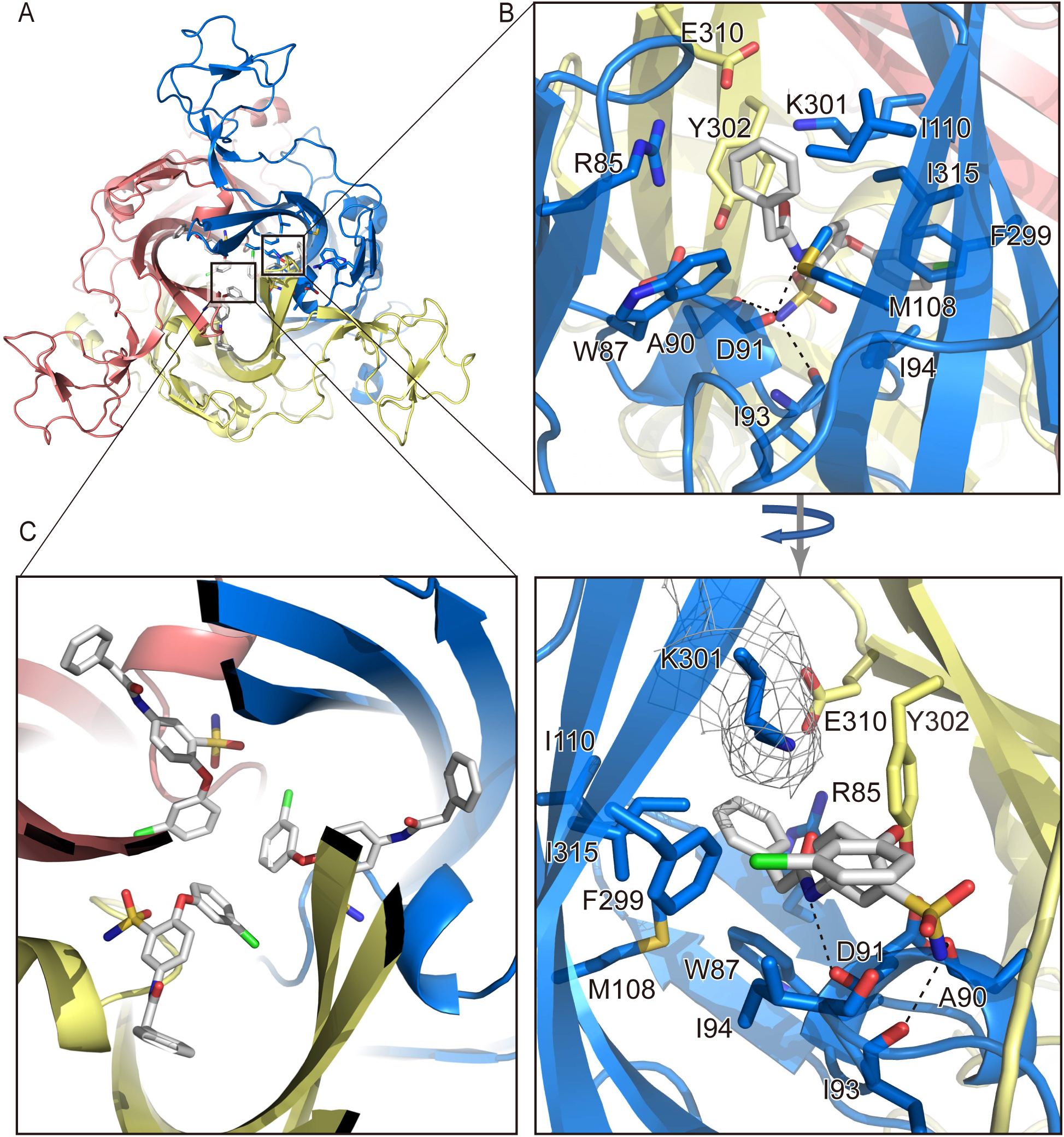
BAY-1797 binding pocket. (**A**) Overall structure of the BAY-1797-bound P2X4 receptor viewed from the extracellular side. The BAY-1797 molecule and amino acid residues involved in its binding in one of three equivalent binding pockets are shown in stick representations. (**B, C**) Close-up views of the BX430 binding pocket. The EM density map for the Lys301 residue is shown and contoured at 3.0 σ (**B**). The BAY-1797 molecules from two other equivalent binding pockets are also shown (**C**). Dotted lines represent hydrogen bonds.

While most of the residues involved in BAY-1797 binding overlap with those involved in BX430 binding, there are also specific interactions in the binding of BAY-1797. The amine of the sulfonamide group forms hydrogen bonds with the main chain carbonyl groups of Ala90 and Ile93 (**Fig. 3B**). Finally, the chlorophenoxy moiety of BAY-1797 protrudes into the center of the receptor and forms hydrophobic contacts with the corresponding portion of the other two BAY-1797 molecules in the trimer (**Fig. 3C**). From another perspective, while the configurations of BX430 and BAY-1797 in the zfP2X4 structure are very similar (**Fig. 2C**), BAY-1797 has bulkier functional groups deeper in the receptor that mediate the BAY-1797-specific interactions (**Fig. 3B, 3C**), which may be related to the lower reported IC_50_ of BAY-1797 compared to BX430.

### Structure-based mutational analysis

To validate the allosteric site in our structures (**Fig. 2, 3, 4A**), we designed and generated a series of mutants of the zebrafish P2X4 receptor (R85A, W87A, I94A, I94W, K301V, K301R, Y302A, and E310K) and performed patch-clamp recording of these mutants expressed in HEK293 cells (**Fig. 2, 4B**). The mutations at Trp87, Ile94, and Lys301 target the residues involved in the direct interactions with the allosteric modulators, whereas the mutations at Arg85, Tyr302, and Glu310 would destabilize the formation of the binding pocket (**Fig. 2, 3**). Consistent with our structures, all mutants showed significantly decreased sensitivity to BX430 compared to the wild-type construct (**Fig. 4B**).

**Figure 4.**
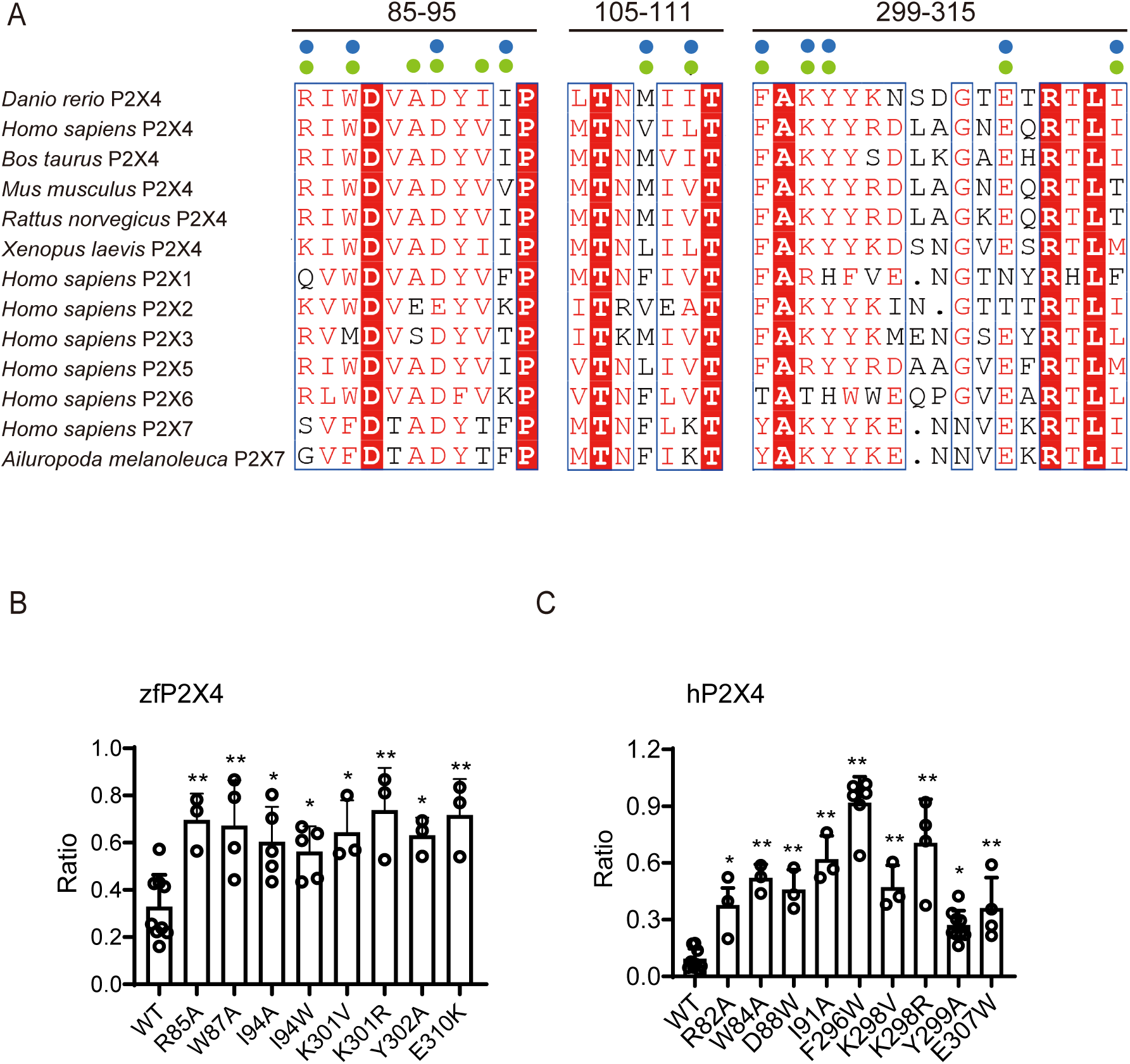
Structure-based mutational analysis. (**A**) Amino acid sequence alignment of P2X4 receptors from *Danio rerio* (zebrafish) (AAH66495.1), *Bos taurus* (NP_001029221.1), *Mus musculus* (NP_035156.2), *Rattus norvegicus* (AAA99777.1), and *Xenopus laevis* (NP_001082067.1), P2X receptors from *Homo sapiens* (P2X1:P51575.1, P2X2:Q9UBL9.1, P2X3:P56373.2, P2X4:Q99571.2, P2X5:Q93086.4, P2X6: O15547.2, and P2X7:Q99572.4), and *Ailuropoda melanoleuca* (giant panda) P2X7 receptor (XP_002913164.3). The residues of the regions located at the binding pocket are shown. Blue and green circles indicate the residues involved in the binding of BX430 and BAY-1797, respectively. **(B)** Effects of BX430 (3 µM) on ATP (1000 µM)-evoked currents of zebrafish wild-type P2X4 and its mutants (mean ± SD, n = 3 to 9). **(C)** Effects of BX430 (5 µM) on ATP (100 µM)-evoked currents of human P2X4 and its mutants (mean ± SD, n = 3 to 9). (ANOVA post hoc test, *: p <0.05, **: p <0.01 vs. WT.)

Next, to verify whether this allosteric site is functionally conserved in the human P2X4 receptor, we also performed patch-clamp recording of hP2X4 and its mutants expressed in HEK293 cells (**Fig. 4C**). The residues involved in the allosteric binding site are highly conserved or at least similarly conserved in both human and zebrafish P2X4 receptors (**Fig. 4A**), and we generated R82A (Arg85 in zfP2X4), W84A (Trp87 in zfP2X4), D88W (Asp91 in zfP2X4), I91A (Ile94 in zfP2X4), F296W (Phe299 in zfP2X4), K298V (Lys301 in zfP2X4), K298R, Y299A (Tyr302 in zfP2X4) and E307W (Glu310 in zfP2X4).

Except for the D88W and F296W mutants (**Fig. 4C**), most of the mutated residues in hP2X4 correspond to the mutated residues in zfP2X4 (**Fig. 4B**). In the BX430-bound structure, the side chain of Asp91 (Asp88 in hP2X4) forms a hydrogen bond with the side chain of Tyr302, contributing to the formation of the binding pocket (**Fig. 2B**). The Phe299 residue (Phe296 in hP2X4) mediates the hydrophobic interaction with BX430 (**Fig. 2B**). In the mutational analysis of hP2X4, all mutants showed significantly reduced sensitivity to BX430 (**Fig. 4C**). Taken together, these results validated the allosteric site of our structures in both zebrafish and human P2X4 receptors.

### Structural reorganization of the binding pocket

Structural comparison with the previously reported apo structure revealed that in the apo state, the intersubunit binding pocket is too narrow to accommodate allosteric modulators (**Fig. 5A, 5B, 5C**). Further close inspection of the allosteric site revealed that in the apo state, the side chain of Lys301 faces the inside of the receptor, which would cause steric hindrance with the pyridine group of BX430 in the BX430-bound structure (**Fig. 5D, 5E**). In contrast, in the BX430-bound and BAY-1797-bound structures, the side chain of Lys301 moves away from the inside of the binding pocket to form a cation-pi interaction with the aromatic ring of Tyr302 from the adjacent subunit and to form hydrophobic interactions with either the 4-isopropylphenyl ring of BX430 or the phenyl ring of the phenylacetamide moiety of BAY-1797 (**Fig. 5F, 5G**, **Supplementary Movie 1)**.

**Figure 5.**
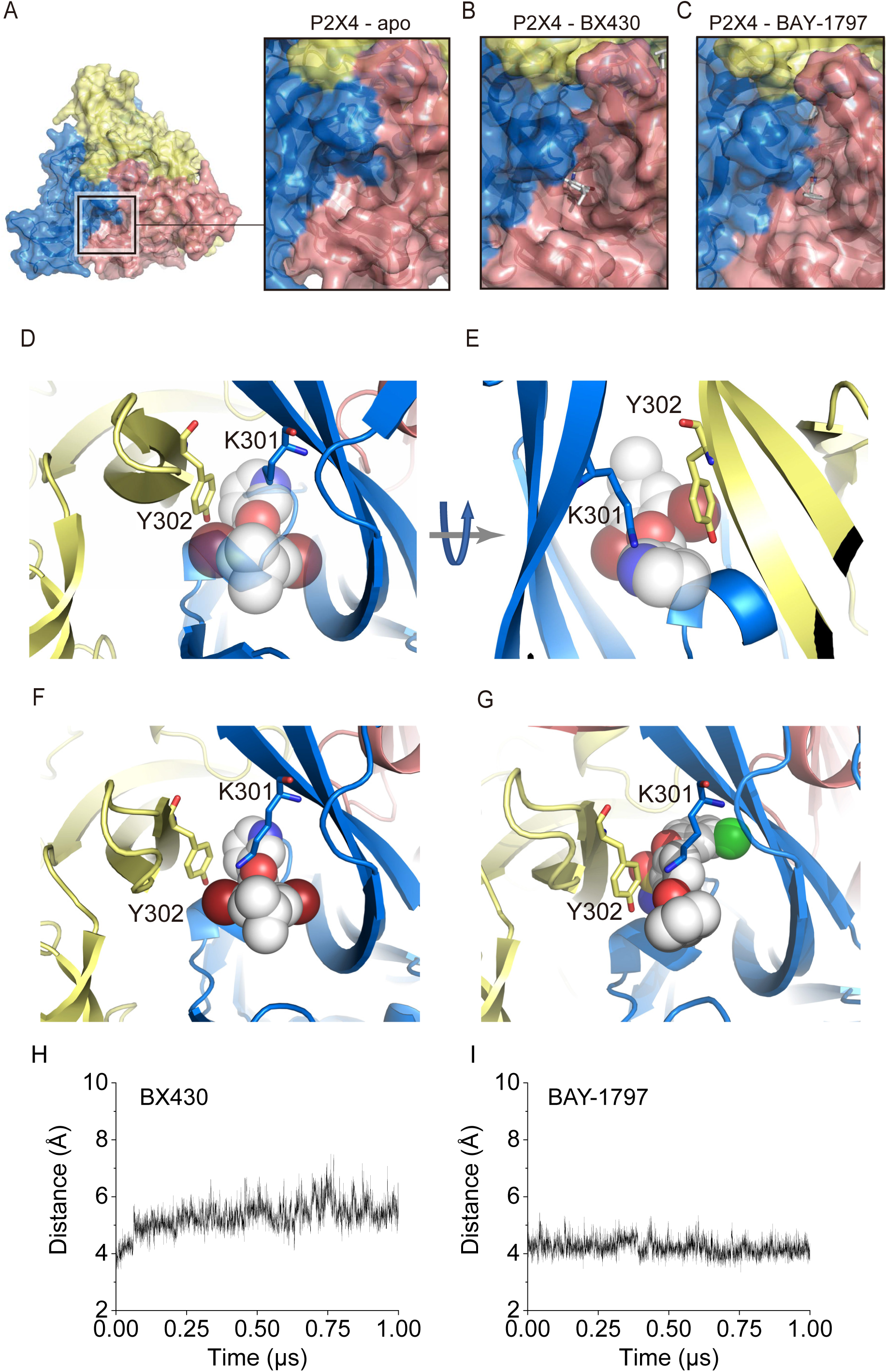
Ligand-dependent structural reorganization of Lys301. (**A-C**) Close-up views of the allosteric site of zfP2X4, viewed from the extracellular side, are shown in surface representations. The zfP2X4 structure in the apo state (PDB ID: 4DW0). (**B**) The BX430-bound zfP2X4 structure (this study). (**C**) The BAY-1797-bound zfP2X4 structure (this study). (**D-G**) in the apo structure (PDB ID: 4DW0) (**D, E**), in the BX430-bound structure (**F**), and in the BAY-1797 bound structure (**G**). The BX430 (**F**) and BAY-1797 (**G**) molecules are shown in sphere representations. The BX430 molecule from the BX430-bound structure superposed onto the apo structure is shown in half-transparent sphere representations (**D**, **E**). **(H, I)** MD simulations of the BX430-bound structure (**H**) and the BAY-1797-bound structure with (**I**). Distance plots between the NZ atom of Lys301 and the center of the aromatic ring of Tyr302. The averaged distance values from the three subunits of the trimer are shown.

In other words, the structural changes in the side chain of Lys301 not only provide enough space at the inner side of the binding pocket to accommodate allosteric modulators (**Fig. 5E**) but also contribute to the direct contacts between Lys301 and allosteric modulators in the outer region of the binding pocket (**Fig. 5F, 5G**). Consistently, the mutations at Lys301 and Tyr302 in zebrafish P2X4 and at the corresponding residues in human P2X4 showed decreased sensitivity to BX430 in patch-clamp recording (**Fig. 4B, 4C**). Furthermore, the intersubunit cation-pi interaction between the side chain of Lys301 and the aromatic ring of Tyr302 was maintained during the MD simulations of the BX430-bound and BAY-1797-bound structures (**Fig. 5H, 5I**).

### Expansion of the binding pocket for allosteric inhibition

In addition to the structural change of Lys301, structural comparison with the previously reported apo structure revealed the expansion of the binding pocket (**Fig. 6A**, **Supplementary Movie 1**). For instance, in the apo structure, the distance between the Cα atoms of Ile110 (chain A) and Gln116 (chain B), located at the entrance region of the binding pocket, is 17.0 Å, whereas the corresponding distances are 18.3 Å and 18.7 Å for the BX430-bound and BAY-1797-bound structures, respectively, reflecting the expansion of the binding pocket (**Fig. 6A**).

**Figure 6.**
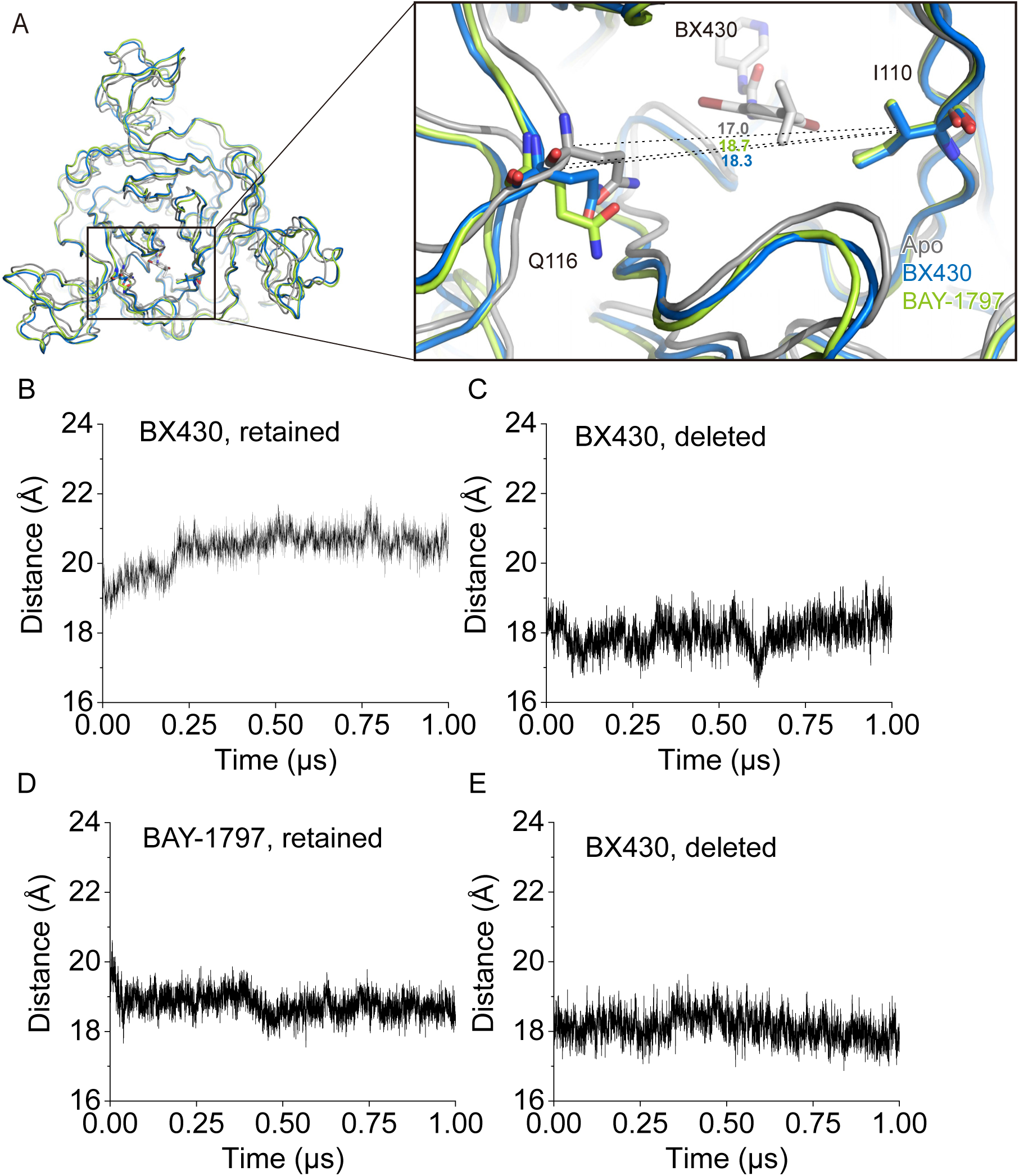
Ligand-dependent expansion of the binding pocket. (**A**) Close-up view of the allosteric site of the zfP2X4 receptor. The BX430-bound (blue) and BAY-1797-bound (green) structures are superposed onto the apo structure (PDB ID: 4DW0, gray). Dotted lines indicate the distance (Å) between the Cα atoms of Ile110 and Gln116 of two adjacent subunits. **(B-E)** MD simulations using the BX430-bound structure with BX430 retained (**B**) or deleted (**C**) and the BAY-1797-bound structure with BAY-1797 retained (**D**) or deleted (**E**) as starting models. The distance plots of Cα atoms between Ile110 and Gln116 of two adjacent subunits are shown. The averaged distances from the three subunits of the trimer are shown. The distance values averaged over the entire 1-µs runs are 20.4 Å (**B**), 18.0 Å (**C**), 18.8 Å (**D**), and 18.1 Å (**E**).

To test whether this expansion is ligand-binding dependent, we performed MD simulations using the BX430-bound structure with BX430 deleted and the BAY-1797-bound structure with BAY-1797 deleted as starting models. In both runs of the MD simulations, both overall structures were mostly stable (**Supplementary Fig. 8B, 8D**). Consistent with our notion, while the binding pocket remains expanded during the MD simulations of the BX430-bound and BAY-1797-bound structures (**Fig. 6B, 6D**), the removal of BX430 and BAY-1797 leads to shorter distances of the Cα atoms of Ile110 and Gln116 of two adjacent subunits of the trimer, indicating a closing motion of the binding pocket in the absence of allosteric modulators (**Fig. 6C, 6E**). It should be noted that the binding pocket is more expanded in the presence of BX430 than in the presence of BAY-1797 (**Fig. 6B, 6D**), probably simply because the 4-isopropylphenyl ring of BX430, located in the outer region of the binding pocket, is bulkier than the phenyl ring of the phenylacetamide moiety of BAY-1797 (**Fig. 5F, 5G**).

As the binding pocket in the body domain expands, the head domain adjacent to the body domain also moves away from the center of the receptor (**Fig. 7A, Supplementary Movie 1**), which is distinct from the downward movement of the head domain upon ATP-dependent channel activation (**Fig. 7B**). For instance, the distance between the Cα atoms of Thr149 in the head domain of the apo- and ATP-bound structures is 0.8 Å, whereas the corresponding distances for the BX430-bound and BAY-1797-bound structures are 2.4 Å and 2.0 Å, respectively, reflecting the movement of the head domain. Furthermore, since the upper body domain acts as a fulcrum in the opening motions of the lower body domain during ATP-dependent activation of P2X4 receptors as well as other P2X subtypes^21, 28, 29^ (**Fig. 7B**), the intersubunit binding of allosteric modulators to the upper body domain would interrupt the structural changes of the body domain (**Fig. 7A**).

**Figure 7.**
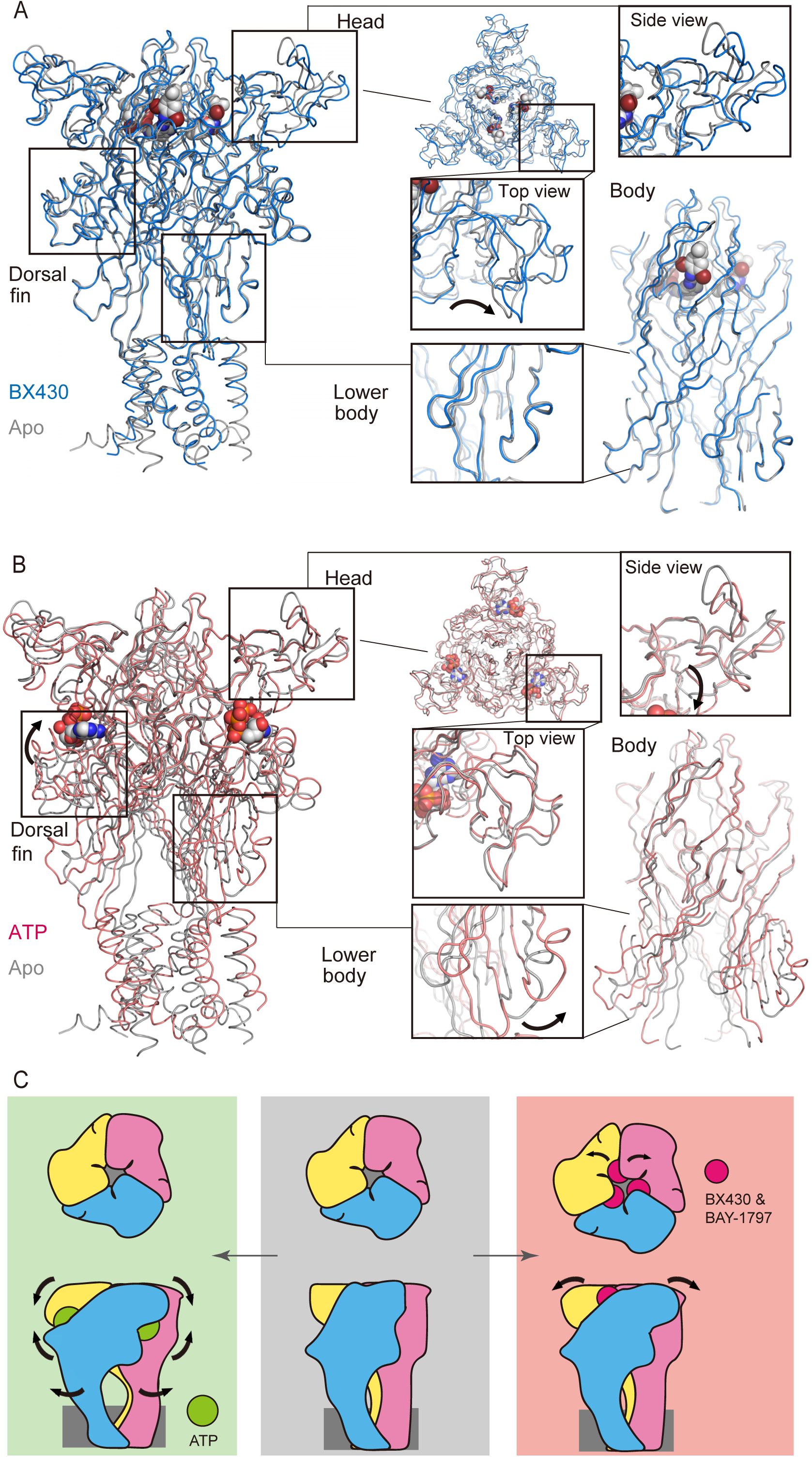
Mechanism of allosteric inhibition. (**A, B**) Superimposition of the BX430-bound (**A**) and ATP-bound (**B**) (PDB ID: 4DW1, red) zfP2X4 structures onto the apo structure. Close-up views of the head and body domains are also shown. Arrows indicate conformational changes. (**C**) Cartoon diagrams of the conformational changes associated with the binding of ATP (left) and allosteric modulators (right). The shape of each subunit follows a dolphin model in **Supplementary Fig. 7.** Arrows indicate structural changes.

In summary, the allosteric modulator-dependent expansion of the binding pocket of the P2X4 receptor induces structural changes in the head domain, as well as conformational stabilization of the upper body domain through the accommodation of modulators, which in turn prevents the structural changes associated with channel activation (**Fig. 7C**).

## Discussion

In this work, we determined the cryo-EM structures of the zebrafish P2X4 receptor in complex with two different negative allosteric modulators, BX430 and BAY-1797 (**Fig. 1**), and performed structure-based mutational analysis of the allosteric site by patch-clamp recording of both zebrafish and human P2X4 receptors (**Figs. 2, 3, 4**). Comparison with the previously reported apo structure revealed the structural rearrangement of Lys301 (**Fig. 5**), the mutation of which decreases sensitivity to BX430 (**Fig. 4**). Further structural comparison and MD simulations revealed the ligand-dependent expansion of the binding pocket (**Fig. 6**), which would be associated with the allosteric inhibition of channel activation (**Fig. 7**).

Interestingly, while the corresponding region of the panda P2X7 receptor, but not of the human P2X3 receptor, was also identified as a binding site for its subtype-specific inhibitors (**Supplementary Fig. 7A, 7C, 7D**), the binding pocket of the panda P2X7 remains open even in the apo state (**Supplementary Fig. 10A, 10B**), unlike that of zfP2X4 (**Fig. 5A, 5B, 5C**). Such a difference between P2X4 and P2X7 is probably because in the apo structure of zfP2X4, the side chain of Lys301 faces the center of the receptor, whereas in the apo structure of panda P2X7, the corresponding region (the side chain of Lys297) is disordered (**Supplementary Fig. 10C**). In the apo structure of zfP2X4, the side chain of Lys301 is stabilized by the side chain of Tyr302 and the main chain carbonyl atom of Ala91 via a water molecule (**Supplementary Fig. 10C**). In contrast, in the panda P2X7 structure, the side chain of Tyr295 would cause a steric hindrance to the formation of such interactions, whereas Tyr295 is replaced by Phe299 in zfP2X4 (**Supplementary Fig. 10C**).

Several previous studies have characterized BX430 by using electrophysiological analysis and *in silico* docking simulations^19, 31, 32^. Notably, two independent groups have used electrophysiology to identify Ile312 in human P2X4 (Ile315 in zebrafish P2X4) as a key residue in the formation of a binding pocket for allosteric inhibition, but their BX430 docking simulation results differ^19, 32^. In an earlier study^19^, the dibromo-isopropylphenyl group of BX430 was oriented toward the inner side of the receptor, which contrasts with our cryo-EM structure (**Fig. 2**). In a more recent report by induced-fit docking^32^, the dibromo-isopropylphenyl group of BX430 was oriented toward the outer side of the receptor. However, in the induced-fit docking, Lys298 in human P2X4 (Lys301 in zebrafish P2X4) did not interact with either BX430 or Tyr299 in human P2X4 (Tyr302 in zebrafish P2X4) but rather pointed away from them, which does not agree with our cryo-EM structure and mutational analysis (**Fig. 4, 5**). In our cryo-EM structure, Lys301 forms a cation-pi interaction with the aromatic ring of Tyr302 and a hydrophobic interaction with the 4-isopropylphenyl ring of BX430 (**Fig. 5**), and the mutations to Lys301 in zebrafish P2X4 and Lys299 in human P2X4 reduced the sensitivity to BX430 (**Fig. 4**). Furthermore, it was also predicted that ligand binding would disrupt the important interaction networks of Asp85, Ala87, Asp88 and Ala297 of human P2X4^32^, which are known to be essential for ATP-dependent activation (Asp88, Ala90, Asp91 and Ala300 in zebrafish P2X4), but this prediction also contrasts with our cryo-EM structure. This discrepancy may also be due to the difference in Lys301, which is adjacent to Asp88, Ala90, Asp91 and Ala300. In addition, another recent report predicted the region in the dorsal fin domain as a potential BX430 binding site, which is also inconsistent with our cryo-EM structure^31^.

Comparison of the allosteric site in our structures with the corresponding region in previously reported P2X structures provided mechanistic insight into the subtype specificity of allosteric modulators for the P2X4 receptor. First, the comparison of the BX430-bound zfP2X4 structure with the previously reported P2X3 structure showed that while most of the residues at the allosteric site are common (**Fig. 4A**), Trp87 and Ile94 in zfP2X4 are replaced by Met75 and Thr82 in human P2X3 (**Fig. 8A**), respectively, which would negatively affect the interactions with BX430. Consistently, the mutations at residues Trp87 and Ile94 in zfP2X4 and the corresponding mutations in hP2X4 caused lower sensitivity to BX430 (**Fig. 4B, 4C**). Next, the comparison of the BX430-bound zfP2X4 structure with the previously reported P2X7 structure showed that Arg85 and Ile110 in zfP2X4 are replaced with the Gly and Lys residues in the panda P2X7 structure, respectively (**Fig. 8B**), which would cause the loss of the salt bridge with the glutamate residue as well as steric hindrance with BX430. In our mutational analysis, the mutation at Arg85 in zfP2X4 and the corresponding mutation in hP2X4 decreased the sensitivity to BX430 (**Fig. 4B, 4C**).

**Figure 8.**
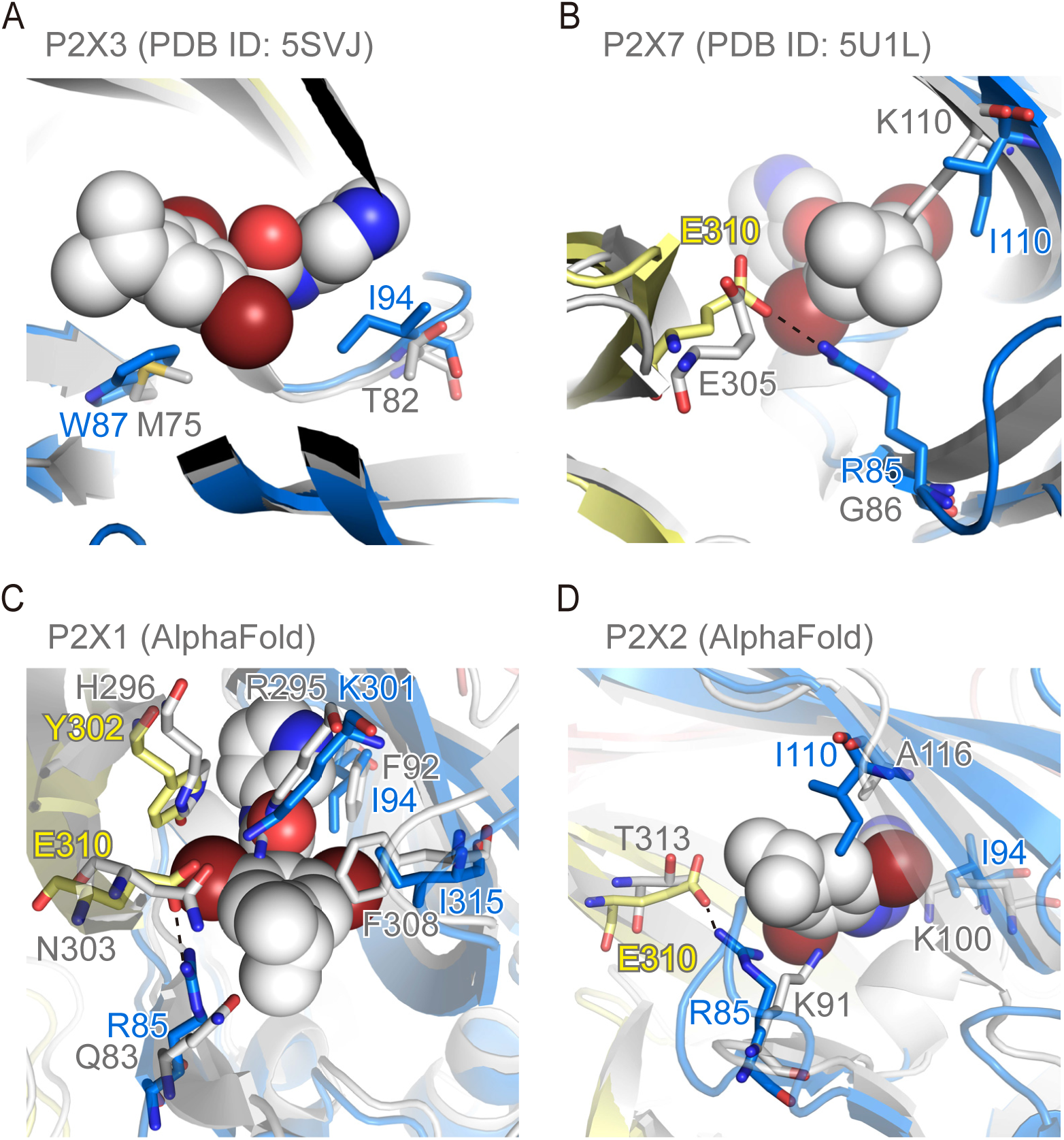
Subtype specificity. (**A-D**) Close-up view of the allosteric site of the BX430-bound zfP2X4 structure. The human P2X3 structure (PDB ID: 5SVJ) (**A**), panda P2X7 structure (PDB ID: 5U1L) (**B**), predicted structure of human P2X1 (AlphaFold) (**C**), predicted structure of human P2X2 (AlphaFold) (**D**) are superposed and shown in gray.

In addition, since no experimental structures have been reported for the other functionally well-characterized P2X receptors (P2X1 and P2X2), we generated predicted structural models of human P2X1 and P2X2 using AlphaFold and ColabFold and superposed them onto the BX430-bound structure of zfP2X4 (**Fig. 8C, 8D**). Arg85, Ile94, Lys301, Tyr302, Glu310, and Ile315 at the allosteric site of zfP2X4 correspond to Gln83, Phe92, Arg295, His296, Asn303, and Phe308 in human P2X1, respectively (**Fig. 8C**), and are shown to be important for BX430 sensitivity either by our mutational analysis (**Fig. 4B, 4C**) or by the previous mutational analysis targeting the residue corresponding to Ile315^19, 32^. In addition, Arg85, Ile94, Ile110 and Glu310 in zfP2X4 are replaced by Lys91, Lys100, Ala116 and Thr313 in human P2X2, respectively (**Fig. 8D**). Among them, the mutations at Arg85, Ile94 and Glu310 in zfP2X4 resulted in decreased sensitivity to BX430 (**Fig. 4B**).

Taken together, our work provides the structural basis for the binding and modulation of the P2X4 receptor by its allosteric modulators, as well as mechanistic insights into their subtype specificity. These insights could contribute to the rational design and optimization of novel allosteric modulators for P2X4 receptors, whose physiological functions are associated with various diseases, particularly neuropathic pain.

## Methods

### Expression and purification

In a previous electrophysiological analysis, the zebrafish P2X4 receptor was shown to exhibit ATP-dependent channel activity even with truncations of 27 residues at the N-terminus and 24 residues at the C-terminus^21^. In this project, we employed an expression construct of zebrafish P2X4 with the less drastic truncation of the N-terminus 8 residues and the C-terminus 7 residues (residues 9-382). The coding region (residues 9-382) of zebrafish P2X4 (GenBank: AAH66495.1) was synthesized (Genewiz, Inc., China) and subcloned into the pFastBac vector with a C-terminal human rhinovirus 3C (HRV 3C) protease cleavage site, coral thermostable GFP (TGP)^33, 34^ and Twin-Strep-tag. DH10Bac *Escherichia coli* cells (Gibco, USA) were used as the host for bacmid recombination. Sf9 cells were cultured in suspension at 27 °C in SIM SF culture medium (Sino Biological, China) and routinely passaged every other day. The initial recombinant baculovirus was generated by transfecting adherent Sf9 cells with the bacmid DNA using the FuGENE HD reagent (Promega, USA) and used to infect suspension cells for virus amplification. The protein was overexpressed in the Sf9 suspension culture at 27 °C for 60 h after virus infection. Cells were collected and lysed by sonication in TBS buffer (50 mM Tris-HCl, pH 8.0, 150 mM NaCl) with 1 mM phenylmethylsulfonyl fluoride (PMSF), 5.2 μg/mL aprotinin, 2 μg/mL leupeptin and 1.4 μg/mL pepstatin A. The supernatant was collected after centrifugation at 8,000 × g for 20 minutes and then ultracentrifuged at 200,000 × g for 1 h. The membrane fraction pellets were solubilized in solubilization buffer (50 mM Tris-HCl, pH 8.0, 150 mM NaCl, 2% n-dodecyl-beta-d-maltopyranoside (DDM) (Anatrace, USA) with 1 mM PMSF, 5.2 µg/mL aprotinin, 2 µg/mL leupeptin, 1.4 µg/mL pepstatin A and 0.2 unit/mL apyrase (Sigma, USA), stirred for 2 h. The solubilized mixture was centrifuged at 200,000 × g for 1 h. The supernatant was loaded onto Strep-Tactin Superflow Plus beads (Qiagen, USA) pre-equilibrated with the wash buffer (100 mM Tris-HCl, pH 8.0, 150 mM NaCl and 0.05% DDM) and stirred for 1 h. The resin was washed with 10 CV of the wash buffer. The proteins were eluted by elution buffer (100 mM Tris-HCl, pH 8.0, 150 mM NaCl, 0.05% DDM, 2.5 mM desthiobiotin) and cleaved by HRV 3C protease to remove TGP and Twin-Strep tag in dialysis buffer (150 mM NaCl, 20 mM HEPES, pH 7.5, 0.05% DDM) overnight. The cleaved protein was applied to a Superdex 200 Increase 10/300 GL column (GE Healthcare, USA) pre-equilibrated with gel filtration buffer (150 mM NaCl, 20 mM HEPES, pH 7.5, 0.05% DDM). Peak fractions were collected and concentrated to ∼2 mg/mL using an Amicon Ultra 50 kDa cutoff (Merck Millipore, USA). All purification steps were performed at 4 °C.

### Amphipol reconstitution

Purified P2X4 was mixed with PMAL-C8 amphipol (Anatrace, USA) dissolved in reconstitution buffer (150 mM NaCl, 20 mM HEPES pH 7.5) at a ratio of 1:10 (w:w) and incubated overnight. Then, detergent was removed after 4 h of incubation with Bio-Beads SM-2 (Bio-Rad, USA) pre-equilibrated with the reconstitution buffer. The sample was applied to a Superdex 200 10/300 GL column (GE Healthcare, USA) pre-equilibrated with the reconstitution buffer. Amphipol-reconstituted P2X4 was pooled and concentrated to approximately 1 mg/mL using an Amicon Ultra 50 kDa cutoff (Merck Millipore, USA). All steps were performed at 4 °C.

### EM data acquisition

For grid preparation, amphipol-reconstituted P2X4 was incubated with 50 μM BX430 or 125 μM BAY-1797 at a final concentration for 1 h. After centrifugation at 100,000 × g for 20 min, the sample was applied to holey carbon-film grids (Quantifoil, Germany, Au R1.2/1.3 μm size/hole space, 300 mesh) glow-discharged for 60 s, blotted with a Vitrobot (ThermoFisher Scientific, USA) with 100% humidity at 4 °C and plunge-frozen in liquid ethane cooled by liquid nitrogen.

Grids were transferred into a Titan Krios (Thermo Fisher Scientific, USA) electron microscope equipped with a K3 direct electron detector (Gatan Inc., USA). Cryo-EM images were collected at an acceleration voltage of 300 kV with a magnification of 29,000× for each dataset. For the dataset of the BX430-bound structure, the pixel size was 0.83 Å, and the defocus was varied from -1.3 μm to -2.0 μm. The dose rate was 20 e^-^/s, and the exposure time was 1.76 s, with an exposure of 50 e^-^/Å^2^. For the dataset of the BAY-1797-bound structure, the pixel size was 0.89 Å, the defocus was varied from -1.4 μm to -2.0 μm, and the dose rate was 18.5 e^-^/s. The exposure time was 1.83 s, with an exposure of 43 e^-^/Å^2^. Detailed data collection statistics are shown in **Table 1**.

**Table 1.** Cryo-EM data collection, refinement and validation statistics

|  | BX430-bound zfP2X4<br>(EMDB-35434)<br>(PDB 8IGJ) | BAY-1797-bound zfP2X4<br>(EMDB-35433)<br>(PDB 8IGH) |
| --- | --- | --- |
| **Data collection and processing** |  |  |
| Magnification | 29000× | 29000× |
| Voltage (kV) | 300 | 300 |
| Electron exposure (e<sup>-</sup>/Å<sup>2</sup>) | 50 | 43 |
| Defocus range (μm) | -1.3 ~ -2.0 | -1.4 ~ -2.0 |
| Pixel size (Å) | 0.83 | 0.89 |
| Symmetry imposed | C3 | C3 |
| Initial particle images (no.) | 2151822 | 2000031 |
| Final particle images (no.) | 101612 | 43282 |
| Map resolution (Å) | 3.4 | 3.8 |
| FSC threshold | 0.143 | 0.143 |
| Map resolution range (Å) | 3.3 ~ 12.3 | 3.6 ~ 12.4 |
| **Refinement** |  |  |
| Initial model used (PDB code) | 4DW0 | 4DW0 |
| Model resolution (Å) | 3.4 | 3.8 |
| FSC threshold | 0.143 | 0.143 |
| Model resolution range (Å) | 3.3 ~ 12.3 | 3.6 ~ 12.4 |
| Map sharpening *B* factor (Å<sup>2</sup>) | -161.1 | -167.2 |
| Model composition |  |  |
| Non-hydrogen atoms | 7480 | 7458 |
| Protein residues | 949 | 948 |
| Ligands | NAG:9, BX4(BX430):3 | NAG:12, BAY(BAY-1797):3 |
| *B* factors (Å<sup>2</sup>) |  |  |
| Protein | 69.7 | 56.4 |
| Ligand | 100.6 | 65.4 |
| R.m.s. deviations |  |  |
| Bond lengths (Å) | 0.002 | 0.03 |
| Bond angles (°) | 0.470 | 0.576 |
| Validation |  |  |
| MolProbity score | 1.30 | 1.62 |
| Clashscore | 5.57 | 8.66 |
| Poor rotamers (%) | 0.38 | 0.39 |
| Ramachandran plot |  |  |
| Favored (%) | 98.09 | 97.13 |
| Allowed (%) | 1.91 | 2.87 |
| Disallowed (%) | 0.0 | 0.0 |

### Cryo-EM data processing

A dataset of 3866 movie stacks of P2X4 with BX430 and a dataset of 4929 movie stacks of P2X4 with BAY-1797 were collected. Motion correction, contrast transfer function (CTF) parameter estimation, particle picking, and further image processing were performed using RELION (version for the dataset of the BX430-bound P2X4 receptor^35^ and version 3.1 for the dataset of the BAY-1797-bound receptor^36^). Particles were extracted with a circle whose diameter was 120 Å, subjected to 2D classification using RELION, and then classified into 8 classes using 3D classification with C1 symmetry imposed. Particles in the best classes were selected, pooled, and classified into 8 classes using 3D classification with C3 symmetry imposed. After refinement and postprocessing, a final map for P2X4 in complex with BX430 at 3.4 Å resolution (using the Fourier shell correlation = 0.143 cutoff criterion) from 101,612 particles was obtained. The resolution of the final map for P2X4 in complex with BAY-1797 was 3.8 Å from 43,282 particles. Local resolutions were estimated using RELION. The workflows for image processing and 3D reconstruction are shown. Figures were created using UCSF Chimera^37^.

### Modeling, refinement, and analysis

Models were built in Coot^38^ using the zebrafish P2X4 receptor structure in the apo state as a template (PDB ID: 4DW0)^39^. Docking of the template model into the cryo-EM maps was performed in PHENIX^40^. The models were manually adjusted in Coot, followed by real-space refinement in PHENIX. Structure figures were generated using PyMOL (https://pymol.org/). The sequence alignment figure was generated using Clustal Omega^41^ and ESPript 3.0^42^. The predicted structural models of human P2X1 and P2X2 were generated using AlphaFold^43^ and ColabFold^44^.

### Electrophysiology

HEK293 cells were cultured in Dulbecco’s modified Eagle medium (DMEM) (Gibco, USA) supplemented with 10% fetal bovine serum (FBS) (PAN-Biotech, Germany), 1% penicillin‒ streptomycin (Gibco, USA), and 1% Glutamax (Gibco, USA) at 37 °C with 5% CO_2_. Plasmids were transfected into HEK293 cells by calcium phosphate transfection 1-3. Nystatin-perforated patch-clamp recordings were performed 24-48 h after transfection at room temperature (25 ± 2 °C) ^45–47^. Patch pipettes were pulled from glass capillaries by a two-stage puller (PC-100, Narishige, Japan), and the resistance was 3-5 megaohms (mΩ) when filled with the pipette solution containing (in mM) 75 K_2_SO_4_, 55 KCl, 5 MgSO_4_ and 10 HEPES adjusted to pH 7.2. The bath solution contained (in mM) 150 NaCl, 0.5 KCl, 10 glucose, 2 CaCl_2_, 10 HEPES and 1 MgCl_2_ adjusted to pH 7.35–7.40. Current traces were filtered at 2 kHz and acquired at 10 kHz via a Digidata 1550B interface and pCLAMP software (Molecular Devices, USA). The membrane potential was held at -60 mV during the recording. ATP or BX430 was dissolved in bath solution and applied to the Y-tube. Between each application, the cells were perfused for 6-8 min to allow for full recovery from desensitization.

### Molecular dynamics simulation

Molecular dynamics simulations were carried out by the program DESMOND^48, 49^. The disordered parts of the TM helices of P2X4 in the cryo-EM structures were manually extended based on the apo state structure of P2X4 (PDB ID:4DW0). Zebrafish P2X4 structures with or without ligands were embedded in a palmitoyl-oleoyl-phosphatidylcholine (POPC) lipid bilayer and dissolved in simple point charge (SPC) water molecules. The systems were neutralized by adding counter ions (Na^+^ or Cl-) to balance the net charges. NaCl (150 mM) was added into the simulation system to represents background salt under physiological conditions. The DESMOND default relaxation protocol was applied to each system prior to the simulation run: (1) 100 ps simulations in the NVT ensemble with Brownian kinetics using a temperature of 10 K with solute heavy atoms constrained; (2) 12 ps simulations in the NVT ensemble using a Berendsen thermostat with a temperature of 10 K and small-time steps with solute heavy atoms constrained; (3) 12 ps simulations in the NPT ensemble using a Berendsen thermostat and barostat for 12 ps simulations at 10 K and 1 atm, with solute heavy atoms constrained; (4) 12 ps simulations in the NPT ensemble using a Berendsen thermostat and barostat at 300 K and 1 atm with solute heavy atoms constrained; (5) 24 ps simulations in the NPT ensemble using a Berendsen thermostat and barostat at 300 K and 1 atm without constraint. After equilibration, the MD simulations were performed for 1.0 μs. Long-range electrostatic interactions were computed using a smooth particle mesh Ewald method. The trajectory recording interval was set to 200 ps, and other default parameters of DESMOND were used during the MD simulation runs. The OPLS-2005 force field was used to model all atoms and their interactions^50, 51^. All simulations were performed on a DELL T7920 graphic workstation with an NVIDA Tesla K40C-GPU. Preparation, analysis, and visualization were performed on a 12-CPU CORE DELL T3610 graphic workstation.

### Statistics and reproducibility

Electrophysiological recordings were repeated 3-8 times. Error bars represent the standard error of the mean. Cryo-EM data collection and refinement statistics are summarized in **Table 1**.

### Data availability

The atomic coordinates and structural factors of the P2X4 structures were deposited in the Protein Data Bank. The accession numbers for the BX430-bound and BAY-1797-bound P2X4 structures are PDB: 8IGJ and EMD: EMDB-35434 and PDB: 8IGH and EMD: EMDB-35433, respectively.

## Acknowledgments

We thank the staff scientists at the Center for Biological Imaging, Institute of Biophysics, and National Center for Protein Science Shanghai (Chinese Academy of Sciences) for technical assistance with cryo-EM data collection (project numbers: CBIapp202007004; 2020-NFPS-PT-005280). We also appreciate the kind support of Prof. Ye Yu (China Pharmaceutical University) as a senior mentor to Prof. Jing Wang. This work was supported by funding from the National Natural Science Foundation of China to M.H. (32071234, 32271244 and 32250610205) and J. W. (32000869). This work was also supported by the Innovative Research Team of High-level Local Universities in Shanghai, a key laboratory program of the Education Commission of Shanghai Municipality (ZDSYS14005), the Open Research Fund of State Key Laboratory of Genetic Engineering, Fudan University (No. SKLGE-2105), and STI2030-Major Projects (No. 2022ZD0207800).

## Author contributions

S.C. expressed and purified P2X4 and performed cryo-EM experiments with assistance from Y.Z., D.S. and X.T. S.C. and M.H. performed model building. W.C. and Y.Z. performed the electrophysiology experiments. J.W. performed the MD simulation. S.C., J.W. and M.H. wrote the manuscript. J.W. and M.H. supervised the research. All authors discussed the manuscript.

## Competing interests

The authors declare no competing interests.

---

**Supplementary Figure 1.**
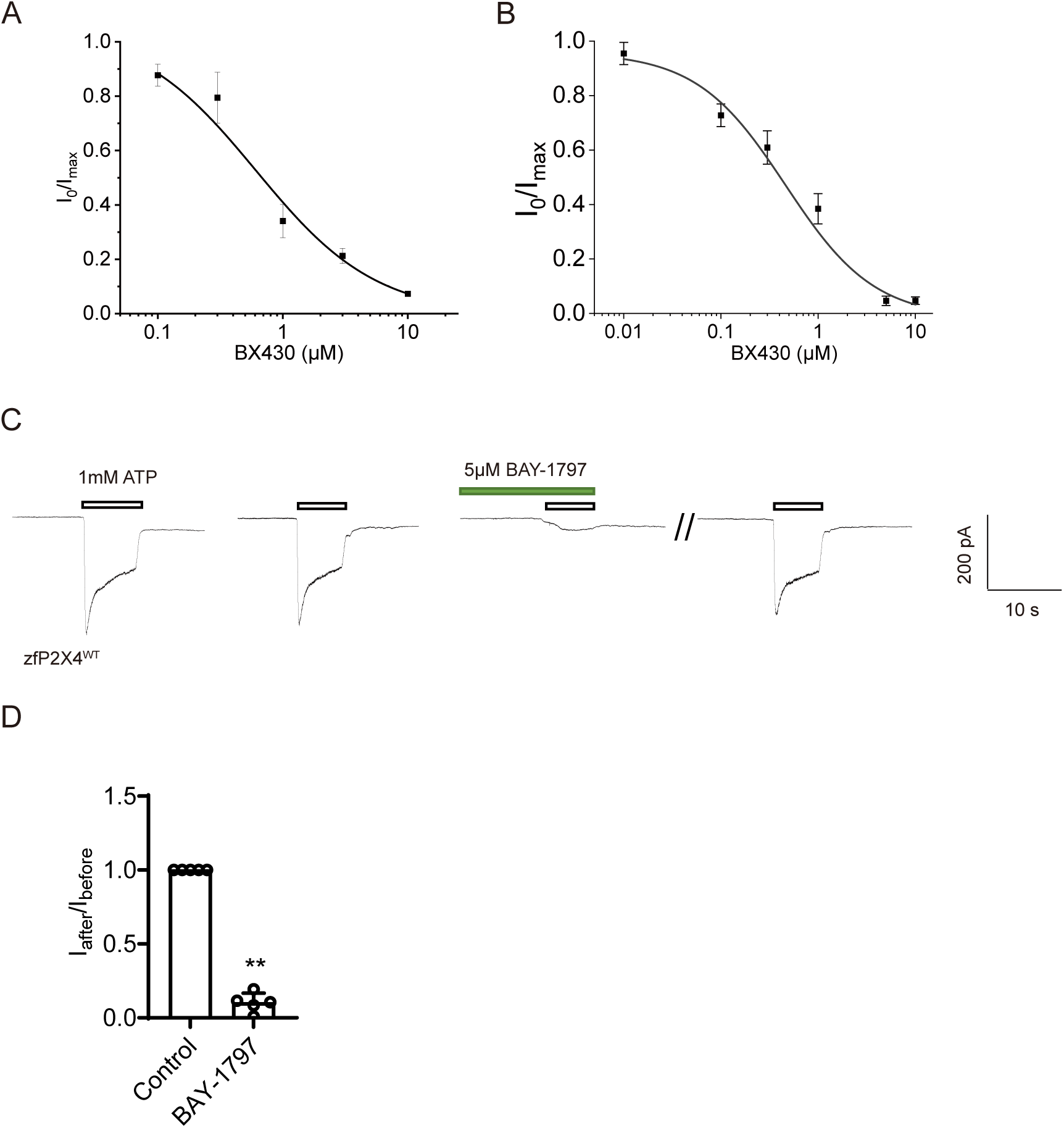
Inhibitory effect of BX430 and BAY-1797 on P2X4 receptors. (A) Concentration‒response curves of BX430 on zebrafish P2X4 (1 mM ATP concentration tested). The estimated IC_50_ value was 0.61 ± 0.66 μM (n=3-4 for each tested BX430 concentration). (B) Concentration‒response curves of BX430 on human P2X4 (100 μM ATP tested). The estimated IC_50_ value was 0.46 ± 0.22 μM (n=7-9 for each tested BX430 concentration). (C) Representative current traces of the effects of BAY-1797 on ATP-evoked currents of zebrafish P2X4. (D) Effects of BAY-1797 (5 µM) on ATP (1 mM)-evoked currents of zebrafish P2X4 (mean ± SD, n = 5) (ANOVA post hoc test, **: p <0.01 vs. WT.).

**Supplementary Figure 2.**
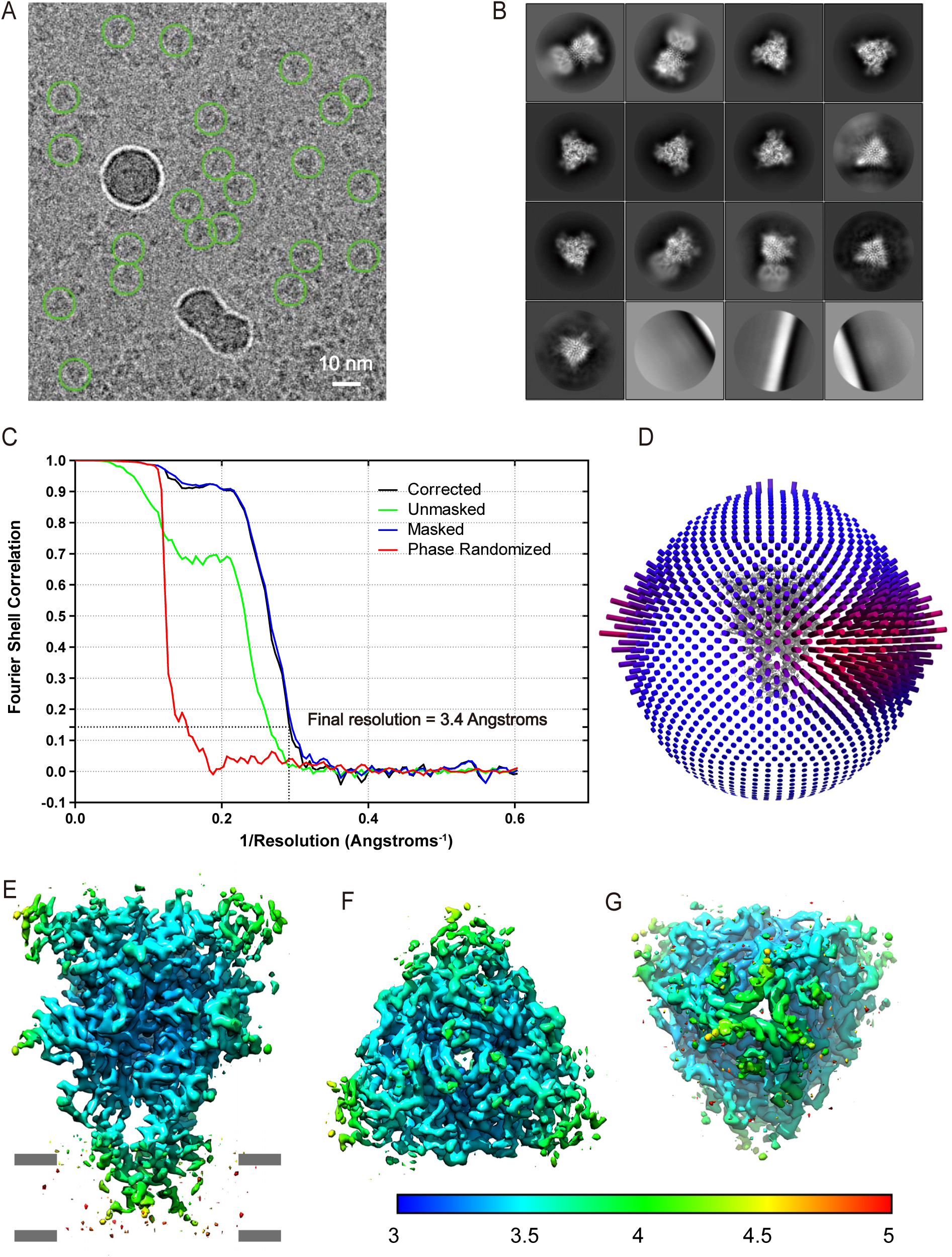
Cryo-EM analysis of zebrafish P2X4 in complex with BX430. (A) Representative cryo-EM image of zebrafish P2X4 particles in complex with BX430. Representative 2D class averages. (C) The gold-standard Fourier shell correlation (FSC) curves for resolution estimation. (D) Angular distribution of the particles used for the final map. (E-G) Side view (E), top view from the extracellular side (F), and bottom view from the cytoplasmic side (G) of the EM density map, colored according to the local resolution, estimated using RELION.

**Supplementary Figure 3.**
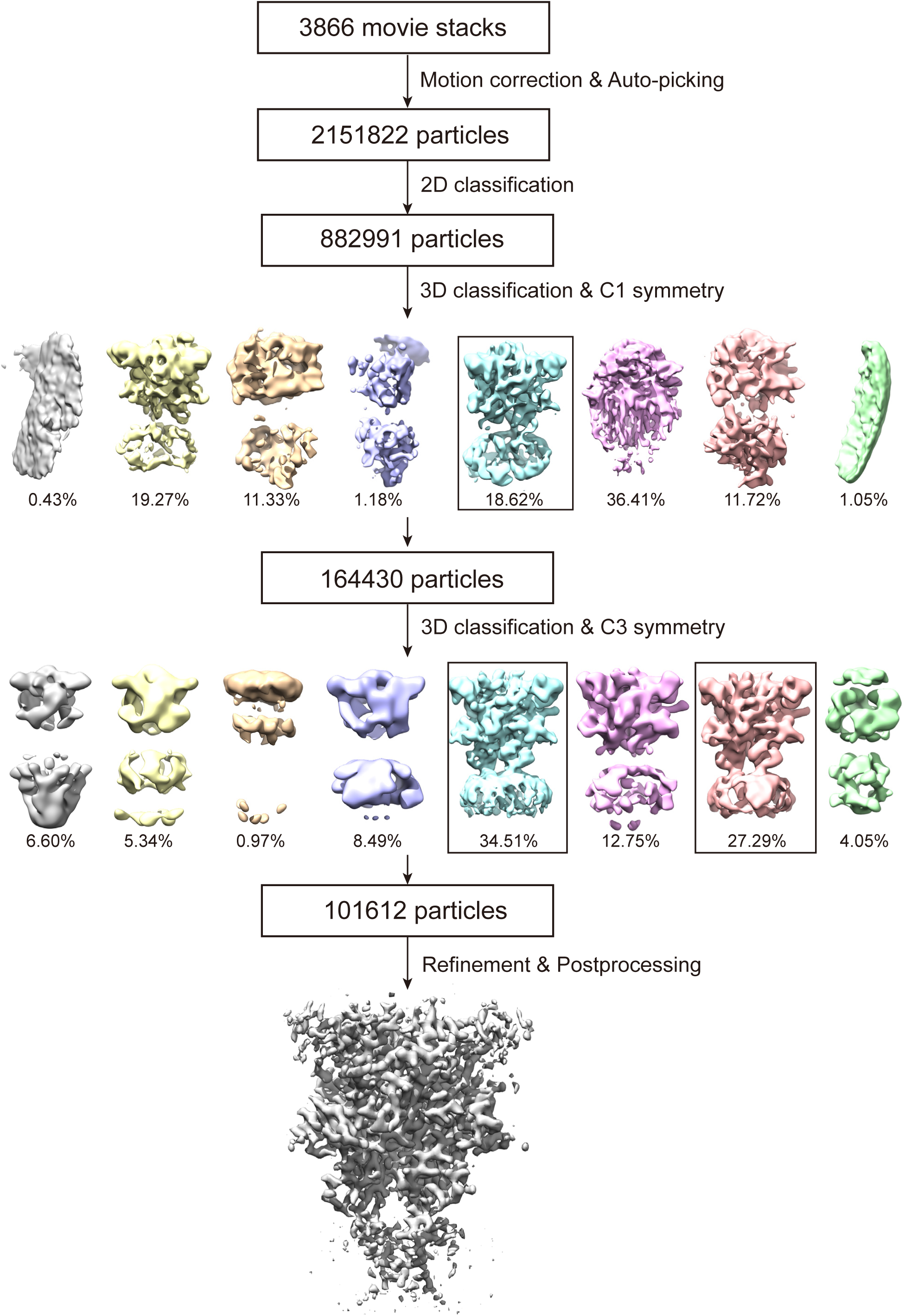
Cryo-EM data processing workflow for zebrafish P2X4 in complex with BX430. All processing steps were performed in RELION. Images were generated using UCSF Chimera.

**Supplementary Figure 4.**
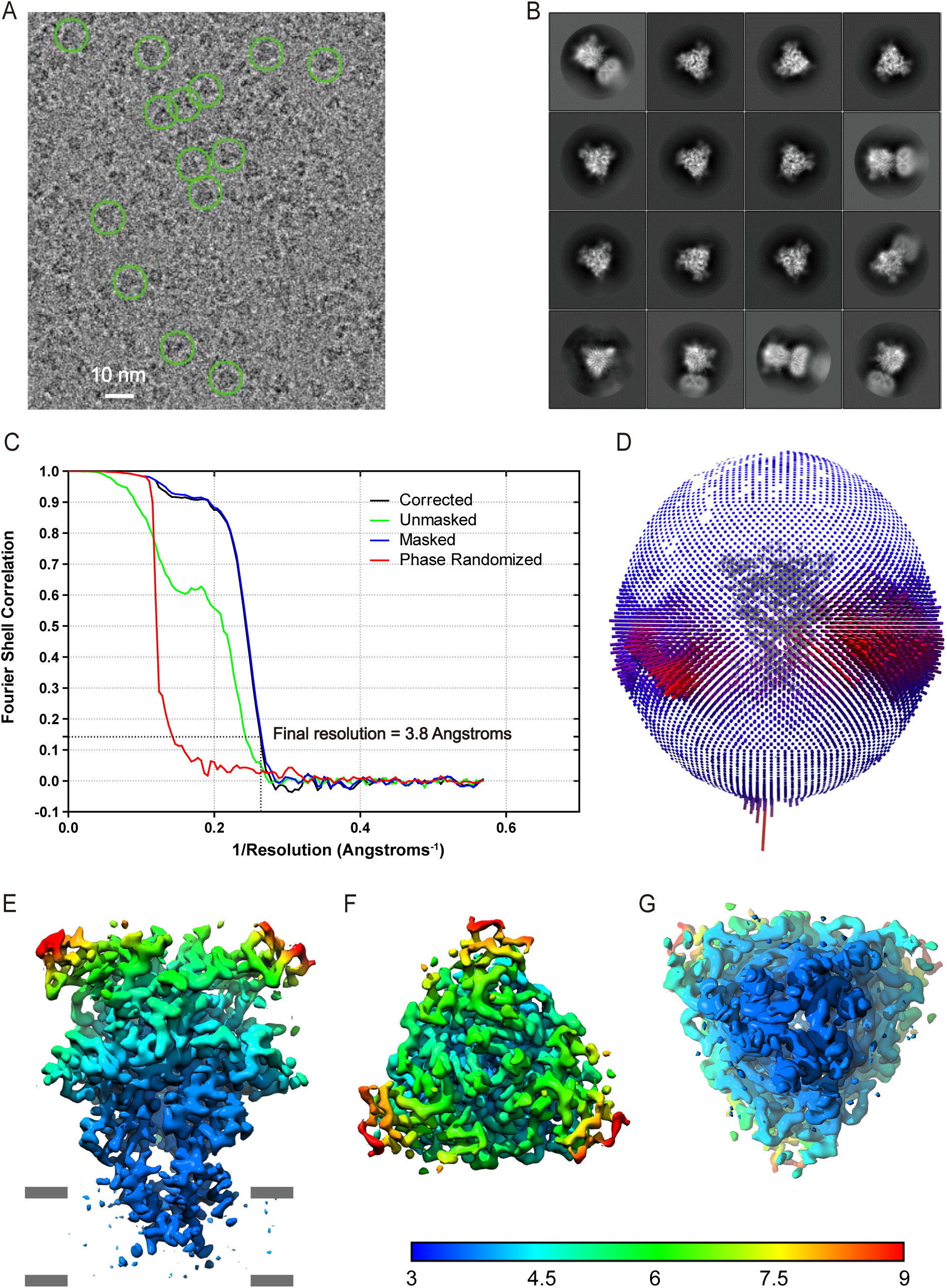
Cryo-EM analysis of zebrafish P2X4 in complex with BAY-1797. (A) Representative cryo-EM image of zebrafish P2X4 particles in complex with BAY-1797. Representative 2D class averages. (C) The gold-standard Fourier shell correlation (FSC) curves for resolution estimation. (D) Angular distribution of the particles used for the final map. (E-G) Side view (E), top view from the extracellular side (F), and bottom view from the cytoplasmic side (G) of the EM density map, colored according to the local resolution, estimated using RELION.

**Supplementary Figure 5.**
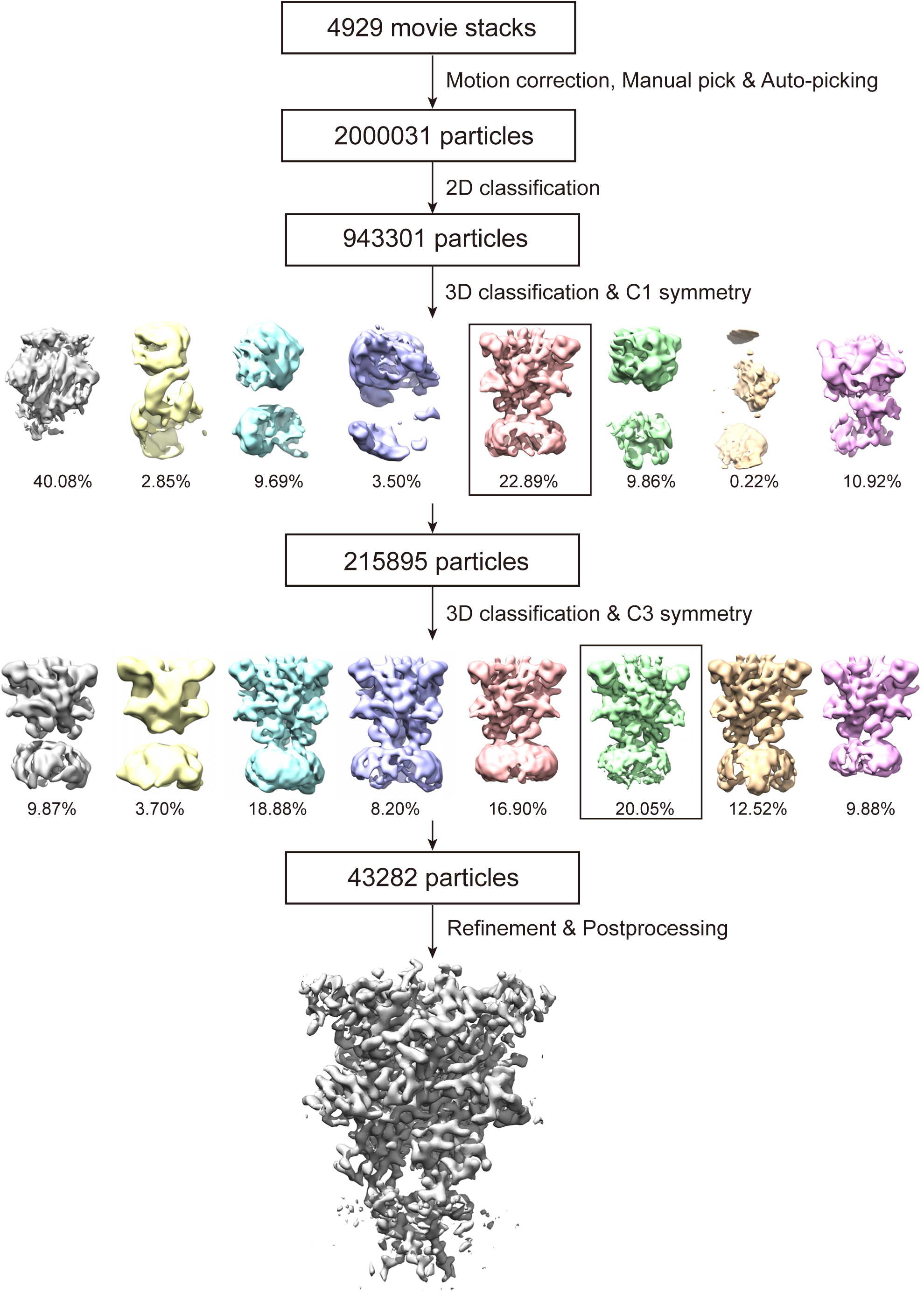
Cryo-EM data processing workflow for zebrafish P2X4 in complex with BAY-1797. All processing steps were performed in RELION. Images were generated using UCSF Chimera.

**Supplementary Figure 6.**
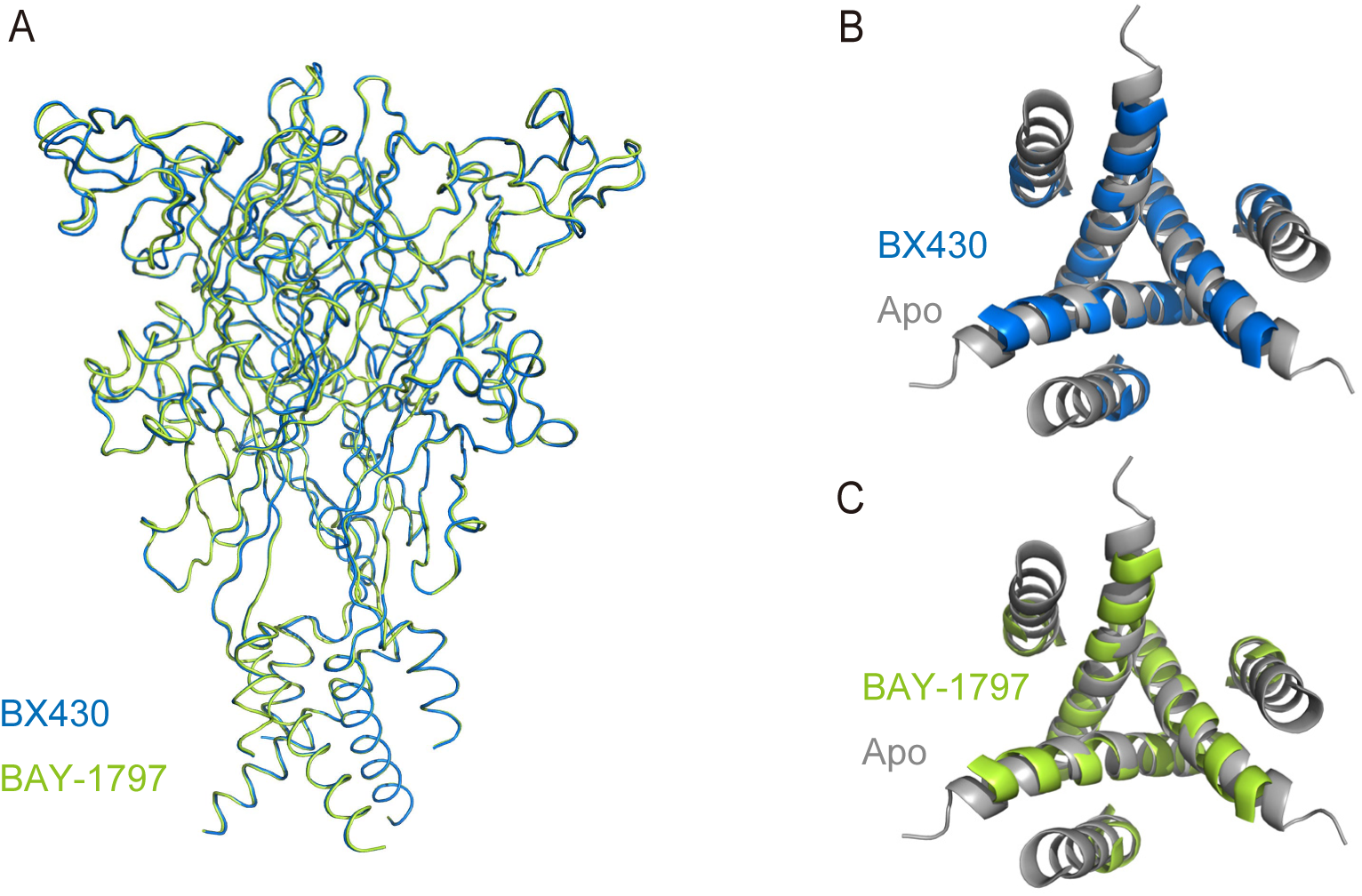
Comparison of the BX430-bound and BAY-1797-bound structures. (A) Superimposition of the BX430-bound structure (blue) onto the BAY-1797 structure (green). (B, C) Superimposition of the BX430-bound (B) and BAY-1797-bound (C) structures onto the apo structure (PDB ID: 4DW0, gray). Only the TM helices are shown and viewed from the intracellular side.

**Supplementary Figure 7.**
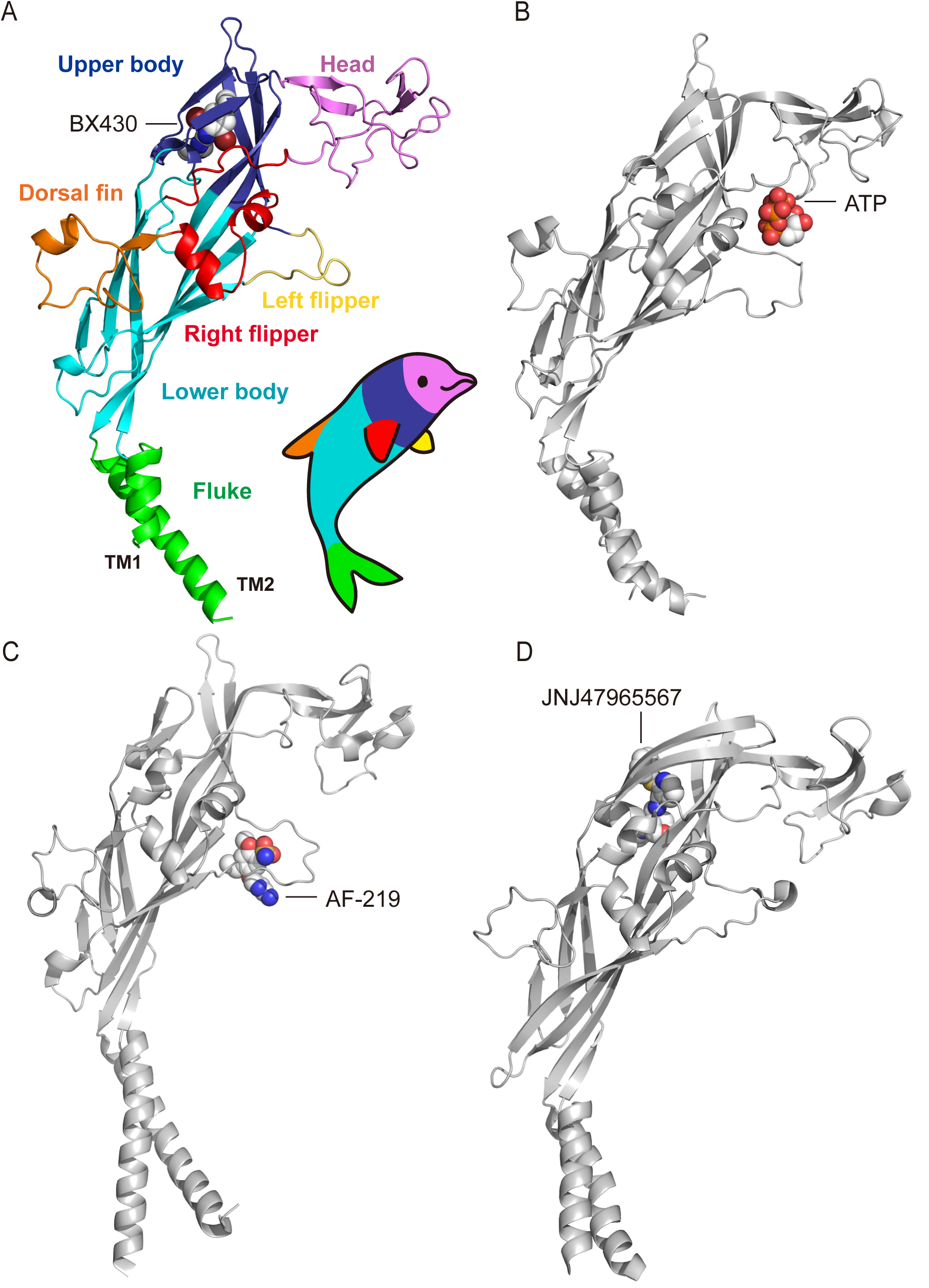
Dolphin model of the P2X subunit. (A) Each region of the P2X4 promoter in cartoon representations is colored according to the dolphin model. (B-D) Each protomer of the ATP-bound P2X4 structure (B) (PDB ID: 4DW1), the AF-219-bound P2X3 structure (C) (PDB ID: 5YVE) and the JNJ47965567-bound P2X7 structure (D) (PDB ID: 5U1X) is shown in gray.

**Supplementary Figure 8.**
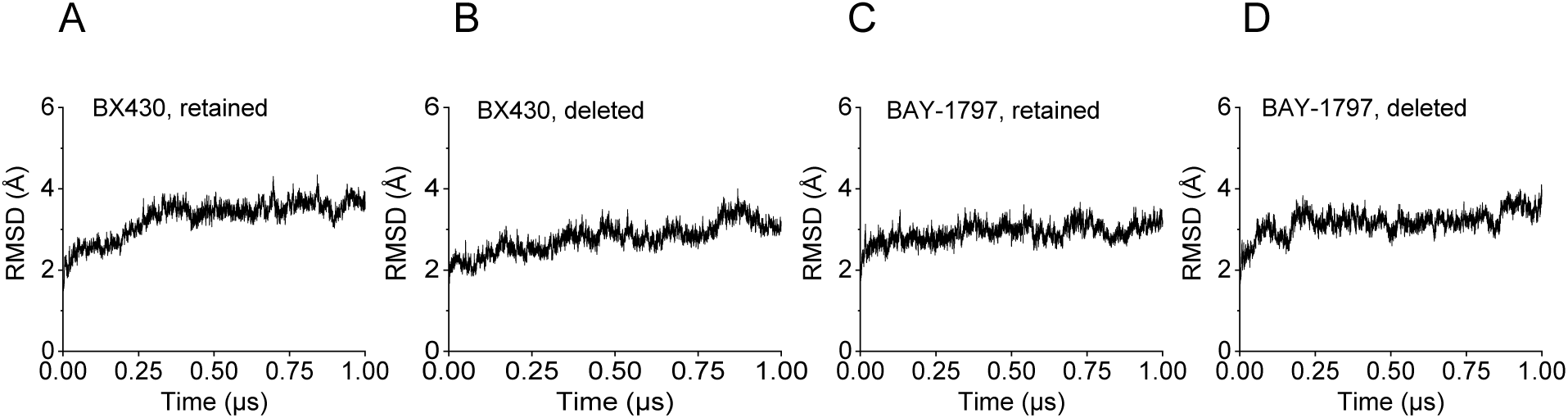
Plots of the RMSD values for Cα atoms during the MD simulations. MD simulations using the BX430-bound structure with BX430 retained (A) or deleted (B) and the BAY-1797-bound structure with BAY-1797 retained (C) or deleted (D) as starting models. The plots of the root mean square deviations (RMSD) for Cα atoms during the MD simulations are shown.

**Supplementary Figure 9.**
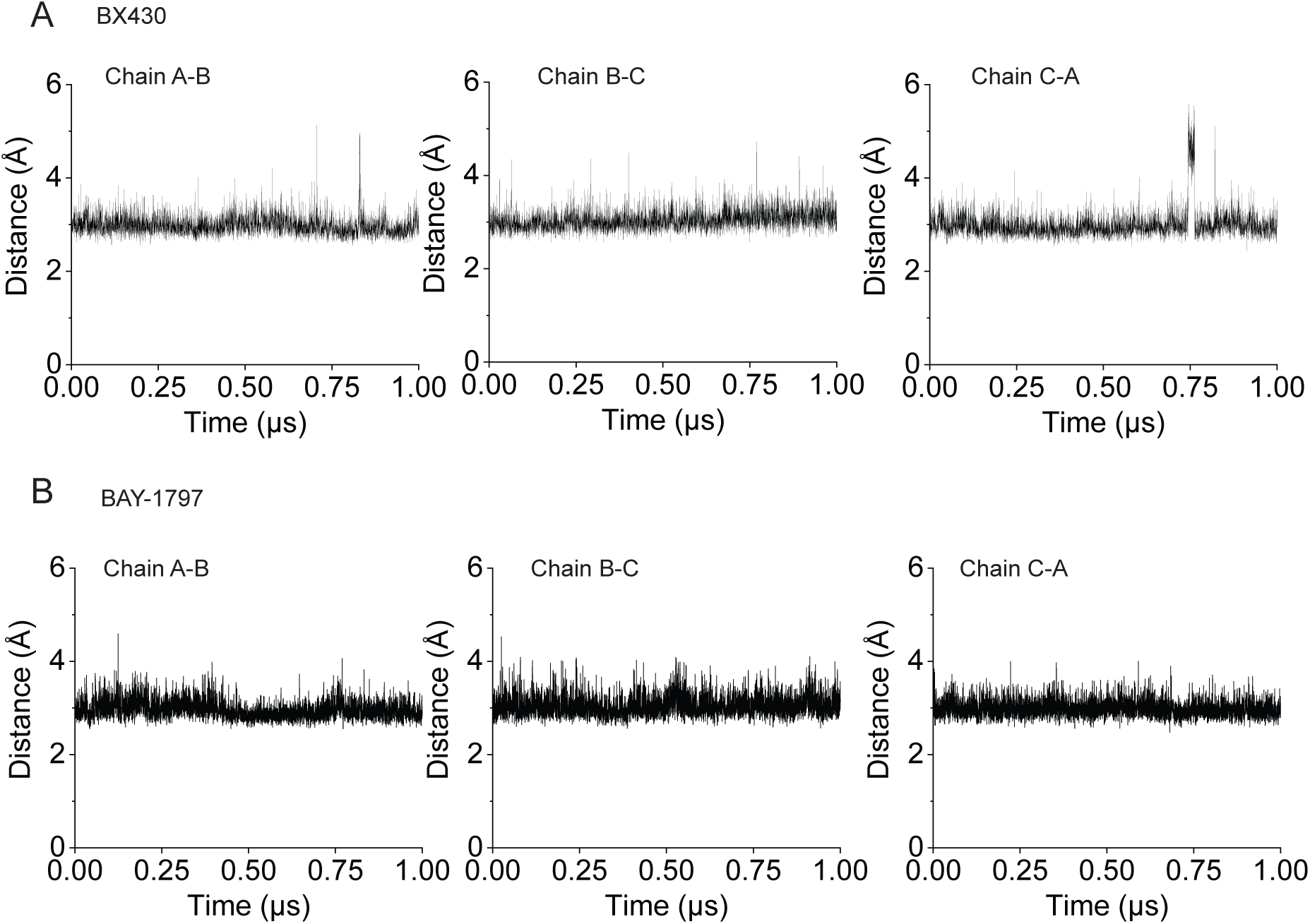
Distance plots between the amine groups of BX430 and BAY-1797 and the main chain carbonyl of Asp91. MD simulations of the BX430-bound structure (A) and of the BAY-1797-bound structure. Plots of the distances between the amine group (N9) of BX430 and the main chain carbonyl of Asp91 of two adjacent subunits (A) and between the amine group (N10) of BAY-1797 and the main chain carbonyl of Asp91 of two adjacent subunits (B) are shown.

**Supplementary Figure 10.**
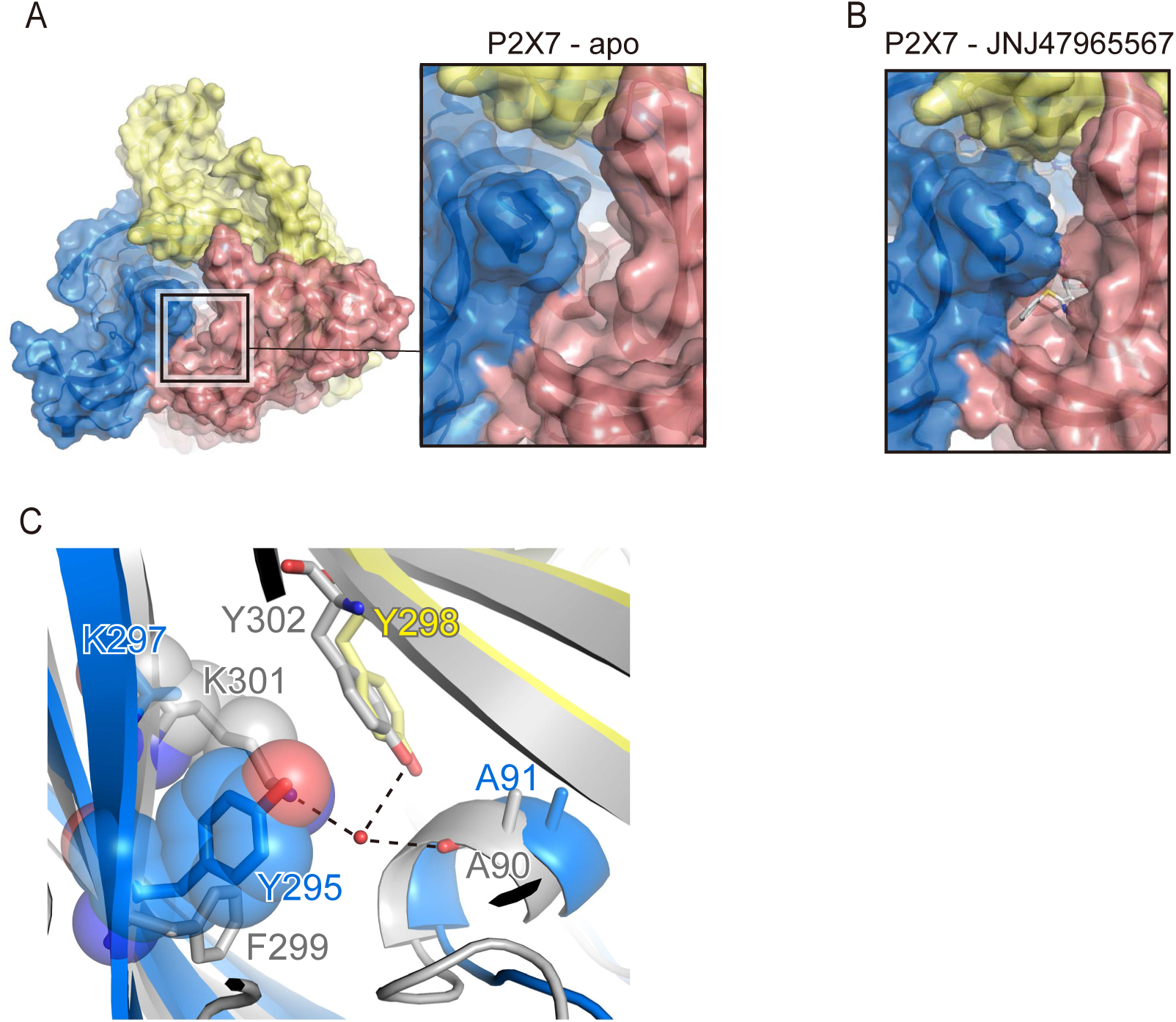
Allosteric sites of P2X7. (A, B) Close-up views of the allosteric site of P2X7, viewed from the extracellular side, are shown in surface representations. (A) The panda P2X7 structure in the apo state (PDB ID: 5U1L). (B) The JNJ47965567-bound panda P2X7 structure (PDB ID: 5U1X). (C) A close-up view of the allosteric site is shown from the panda P2X7 structure in the apo state (colored) superposed onto the zfP2X4 structure in the apo state (gray). Tyr295 in panda P2X7 and Lys301 in zfP2X4 are shown in half-transparent sphere representations. Water molecules are shown as red spheres. Dotted lines indicate hydrogen bonds.

## Notes

### Competing Interest Statement

The authors have declared no competing interest.

